# Molecular insights into bulk lipid transport from structural studies of the bridge-like protein VPS13A complexed with the scramblase XKR1

**DOI:** 10.64898/2026.01.07.698282

**Authors:** Bodan Hu, Daniel Álvarez, Cristian Rocha-Roa, Valentin Guyard, Dazhi Li, Xinbo Wang, Pietro De Camilli, Stefano Vanni, Karin M. Reinisch

## Abstract

In eukaryotes, bridge-like lipid-transfer proteins (BLTPs) are central in mediating vesicle-independent lipid transfer between organelles. BLTPs span the cytosolic space between organelles at contact sites, featuring hydrophobic channels for lipids to travel between membranes. How BLTPs cooperate with partner proteins to orchestrate lipid delivery remains mysterious. Here we used cryo-electron microscopy to visualize a complex comprising the prototypical BLTP VPS13A and the plasma membrane localized scramblase XKR1 at near-atomic resolution. VPS13A interacts with XKR1 via its PH-domain, priming VPS13A’s bridge-like lipid-transfer domain to deliver lipids directly to the cytosolic leaflet of the acceptor membrane. In molecular dynamics simulations, such arrangement allows for robust lipid transfer, accelerated by membrane properties. Newly delivered lipids can then be equilibrated between leaflets of the membrane bilayer by the scramblase, allowing for membrane growth. Mechanistic insights regarding lipid delivery by VPS13A are directly applicable to all VPS13 proteins and all BLTP family members more broadly.

## Introduction

In eukaryotes, the lipids that constitute organellar membranes are synthesized primarily in the endoplasmic reticulum (ER), before being redistributed to other organelles. While long known that lipid transfer between intracellular membranes can take place via vesicular trafficking, it is recently appreciated that bridge-like lipid transfer proteins (BLTPs) play a central and fundamental role in bulk lipid movement, independent of vesicles. BLTPs span between organelles at sites of close apposition via a “bridge-domain”, featuring a hydrophobic groove along which lipids can flow between organellar membranes (Reinisch et al., 2025).

BLTP-mediated bulk lipid transfer is now believed to be key in membrane expansion accompanying the biogenesis of new organelles like the autophagosome (Melia, 2023), membrane maintenance for organelles outside vesicle trafficking pathways like mitochondria (Reinisch and Prinz, 2021), or membrane repair, for example of damaged lysosomal membranes (Wang et al., 2025b). The founding members of the BLTP family, the VPS13 proteins, are also of great biomedical interest because their dysfunctions underlie neurological diseases, such as Parkinson’s (VPS13C) or chorea acanthocytosis (VPS13A); and they, along with the ATG2 family that functions in autophagy, are the most studied BLTPs (Hanna et al., 2023; Reinisch et al., 2025).

An emerging paradigm, based primarily on studies of VPS13 and ATG2 proteins (Adlakha et al., 2022; Ghanbarpour et al., 2021; Maeda et al., 2020; Matoba et al., 2020), is that at the acceptor membrane BLTPs cooperate with membrane-embedded scramblases, proteins that equilibrate phospholipids between membrane leaflets, allowing for the concomitant expansion of both leaflets of the membrane even as the BLTP delivers lipids only to one of the leaflets. The mechanisms governing the transfer of lipid from the bridge-domain to the membrane and the role of the scramblase in this process have been elusive in the absence of near-atomic resolution information regarding BLTP-scramblase complexes.

Here we combine structural and computational approaches to interrogate how a member of the VPS13 family, human VPS13A, partners with a plasma membrane scramblase, XKR1, for lipid delivery. Loss-of function mutations in VPS13A result in chorea-acanthocytosis, a Huntington-like neurodegenerative condition accompanied by abnormal red cells in the blood (acanthocytes)(Rampoldi et al., 2001). A very similar condition, called McLeod’s syndrome, is due to loss of function mutations in XKR1 (Ho et al., 1994), consistent with a functional partnership between the two proteins and their reported direct interaction (Park and Neiman, 2020; Ryoden et al., 2022). Moreover, VPS13A and XKR1 were top hits in a screen for genes required for surface exposure of phosphatidyl serine in T cell (Ryoden et al., 2022), suggesting a model according to which VPS13A delivers phospholipids from the ER to the plasma membrane, where XKR1 scrambles them (Guillen-Samander et al., 2022). VPS13A also localizes to other sites in the cell, such as contacts between the ER and mitochondria, where its function is less well investigated (Kumar et al., 2018).

We used cryo-electron microscopy to visualize the complex of VPS13A, a recently identified soluble interaction partner calmodulin (CaM) (Li et al., 2025), and detergent solubilized XKR1 at near atomic resolution (3.1-3.5 Å, in four different maps). The interaction between VPS13A and XKR1 orients the VPS13A bridge domain in direct vicinity of the lipid bilayer, and pins VPS13A close to the scramblase so that lipids can be scrambled as soon as they are delivered. Molecular dynamics simulations show that the cryo-EM structure is compatible with robust direct lipid transfer between the bridge domain and the cytosolic leaflet of the acceptor membrane, and suggest that membrane properties, namely membrane tension, might couple lipid delivery with lipid scrambling by XKR1. We expect that the mechanistic insights regarding lipid delivery by VPS13A described here are directly applicable to all VPS13 and ATG2 proteins as well as, more broadly, other BLTP families.

## Results

### VPS13A/CaM-XKR1 complex preparation, cryo-EM single particle reconstruction, modelling

We separately produced FLAG-tagged human XKR1 (3XFLAG-XKR1) and VPS13A (VPS13A-3XFLAG) by transfection into Expi293 cells. XKR1 was isolated from the glyco-diosgenin (GDN)-solubilized membrane fraction by affinity purification via anti-FLAG-resin; VPS13A was similarly made, co-purifying with a recently identified interaction partner CaM (Li et al., 2025), from the soluble fraction in the absence of detergent. XKR1 and VPS13A/CaM were mixed together, and the ensuing complex was isolated by size exclusion chromatography, then subjected to cryo-EM single particle analysis (Sup. Fig. S2). We generated two maps, one for VPS13A/CaM and another centered on XKR1 and the VPS13A PH domain. Local refinement yielded a series of maps with nominal resolutions ranging from 3.2-3.5 Å (Fig. 1, Sup. Fig. S2). The maps allowed us to model most of the VPS13A/CaM-XKR1 complex at near-atomic resolution. The portions of VPS13A visualized include most of the bridge-domain as well as C-terminal adaptor modules responsible for its attachment to acceptor membranes: the VAB (poorly resolved), WWE (poorly resolved), and PH “adaptor” domains (Fig. 1). Density for CaM was also well defined. Amphipathic helices at the membrane-proximal end of the bridge-domain, conserved across all VPS13’s and ATG2 proteins (and referred to as the ATG2_C region for their location at the C-terminus of ATG2 where they are also present, Fig. 1A), are not visible in the reconstruction and they were not modeled. The taco-shell shaped bridge domain comprises a series of ten repeating-beta-groove (RBG) modules (Levine, 2022) arranged end to end, sandwiched between N- and C-terminal “caps”. Each RBG module consists of five highly curved antiparallel beta strands, whose concave surface is lined entirely by hydrophobic residues and so suited to solubilize lipid fatty acid moieties, and connecting segments. To model the bridge-domain we segmented an AlphaFold-generated model (Jumper et al., 2021) of the bridge-domain into RBG modules and rigid-body fitted the RBG fragments into the experimental maps. We similarly placed AlphaFold-generated coordinates for XKR1 and adaptor domains of VPS13A (VAB-,

**Figure 1.**
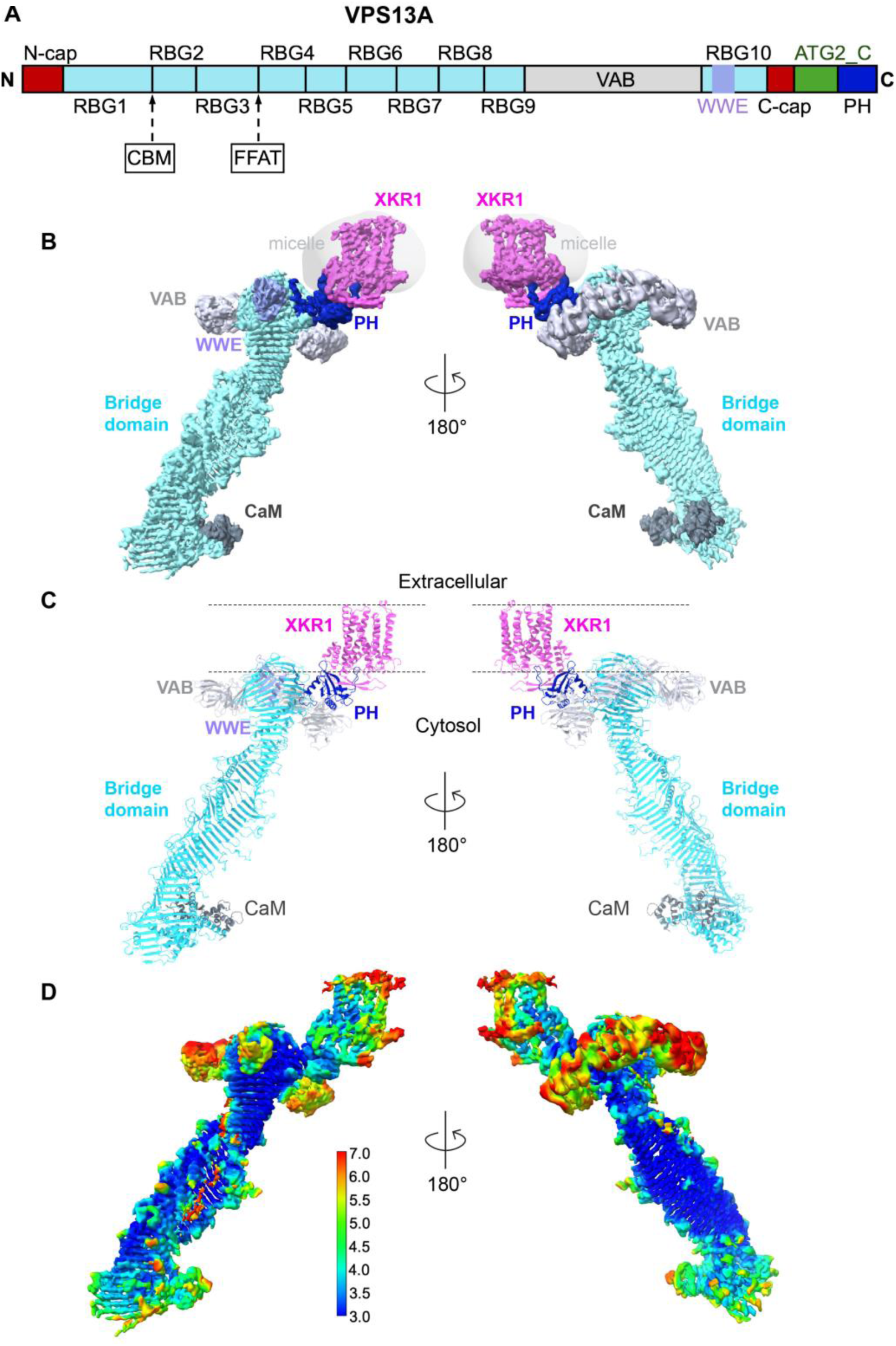
Architecture of the VPS13A/CaM-XKR1 complex. **(A**) Schematic of VPS13A domain architecture. CBM, calmodulin binding motif; FFAT, peptide motif involved in VPS13A-ER association. (**B)**, cryo-EM map of the VPS13A/CaM-XKR1 complex, with the membrane protein XKR1 embedded in GDN micelle. Color coding for VPS13A is as in (A); XKR1 is pink, and CaM is slate gray. **(C)**, Model of the VPS13A/CaM-XKR1 complex, shown in the same color and orientation as the map in (B). **(D)**, Local resolution of cryo-EM map of the VPS13A/CaM-XKR1 complex, shown in the same orientation as in (B).

WWE-, and PH-domains) and CaM into the maps. Once the initial model was assembled, we combined artificial intelligence based map interpretation in DeepMainMast, rigid body fitting in ChimeraX, and manual rebuilding in COOT to construct a final model (Emsley et al., 2010; Meng et al., 2023; Terashi et al., 2024). Except for the VAB and the bridge-domain “caps” in the lowest resolution parts of the reconstruction, the model was subjected to real space refinement in Phenix (Liebschner et al., 2019). (See Sup. Table S1 for data collection and model statistics, and Sup. Fig. S3 for representative density.)

### Overview of the complex

At low resolution, VPS13A resembles a bubble wand (Fig. 1 B-D). It comprises the elongated bridge domain serving as the bubble wand’s “handle”, and emanating from it, the C-shaped VAB, which is prominent as the largest of the adaptor domains and corresponds to the bubble wand’s “loop”. The VAB and the smaller PH and WWE domains are arranged in a plane encircling the C-terminal end of VPS13A’s bridge domain, all positioned to interact with the acceptor membrane and/or membrane embedded receptors like XKR1. CaM binds to the N-terminal end of the bridge domain.

VPS13A connects to the detergent-micelle embedded XKR1 via its C-terminal end. The WWE and VAB adaptor domains are not required for VPS13A’s interaction with XKR1 or VPS13A’s recruitment to the plasma membrane (Guillen-Samander et al., 2022), and accordingly they do not contact XKR1. VPS13A interacts with detergent solubilized XKR1 solely via its PH adaptor domain, which protrudes rigidly from the back of the taco shell-shaped bridge domain (Fig. 1 B-C). The PH domain is positioned over XKR1 in the membrane and does not appear to access membrane directly, except that two loops—PH-L_ß3-ß4_ and PH-L_ß6-ß7_—independently interact with residues of XKR1 transmembrane helices (see below).

Importantly, there is no direct interaction between the bridge-domain of VPS13A and XKR1; and the bridge domain, whose long axis is tilted ∼35° from a normal to the membrane plane as defined by the micelle surface, is positioned to deliver lipids directly to the acceptor membrane (Fig. 1 B-C).

### VPS13A bridge domain and bound lipids

As expected from AlphaFold predictions, the bridge domain forms a ∼240 Å long extended beta sheet with a taco shell shaped structure, whose interior is lined by hydrophobic residues to form the lipid transport groove and whose exterior is hydrophilic (Fig. 2). In the experimental structure, the taco shell twists by 270° over its length. The width and depth of the lipid transfer groove vary along the length of VPS13A. The width across the groove ranges from ∼25 Å across at its widest points near the ends, to ∼9 Å across at its narrowest points (Fig. 2 A-B), just wide enough across to accommodate 1-2 lipids. The width across other experimentally characterized VPS13s (VPS13C and fungal VPS13, (Li et al., 2025) & (Li et al., 2020)) and BLTPs (ATG2, BLTP1, (Kang et al., 2025; Wang et al., 2024)) is similarly variable. Whether the variability is relevant for the transfer mechanism is not known, although it is intriguing to speculate that modulating channel width could regulate lipid flow rates.

**Figure 2.**
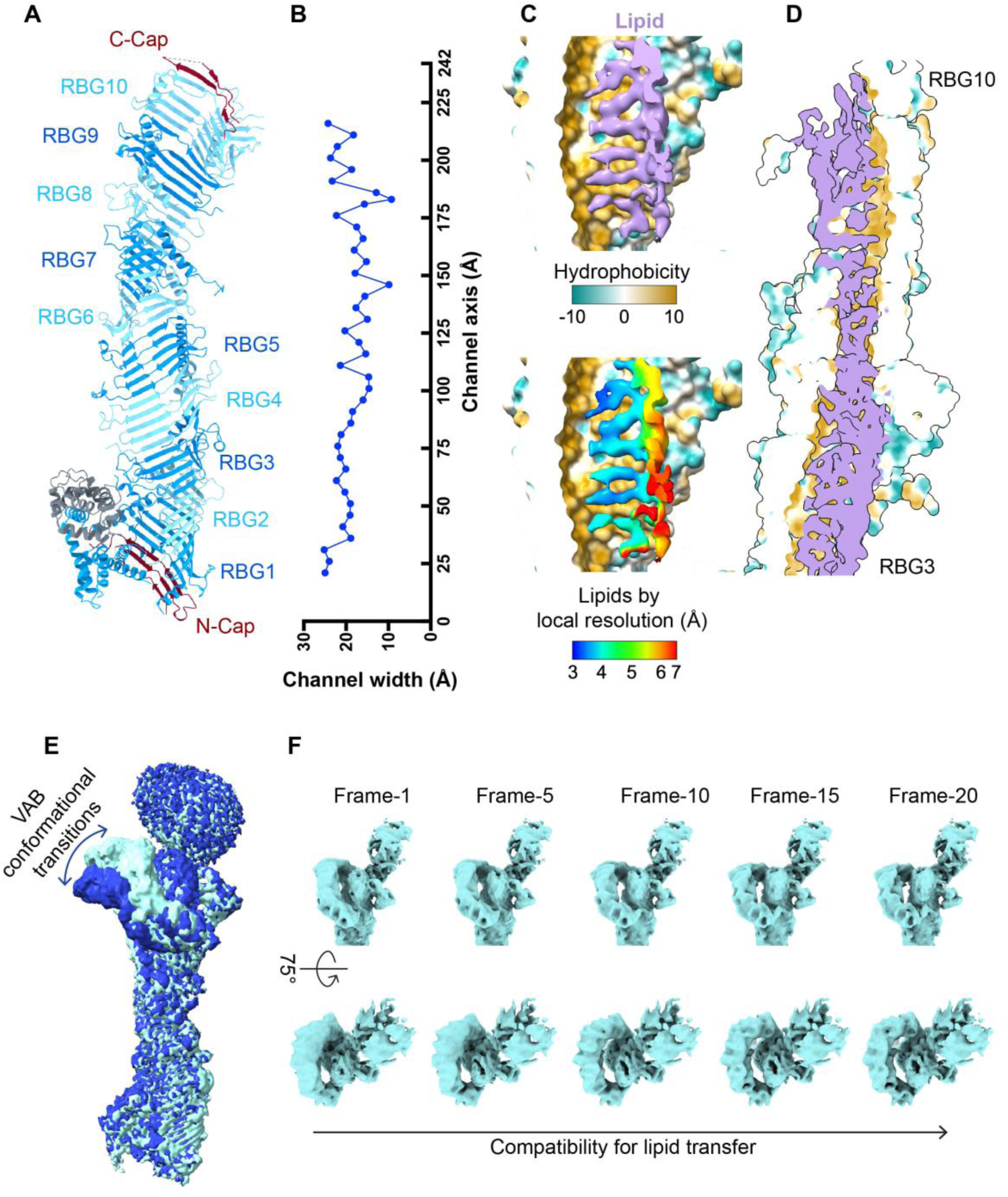
Structure, lipid binding, and conformational dynamics of the VPS13A bridge domain. **(A)** The bridge domain comprises ten repeating β-groove (RBG) motifs and modeled portions of the cap regions located at the N-and C-termini of the lipid transfer channel. **(B)** Width of the lipid transfer channel calculated using ChimeraX, excluding the N-terminal and C-terminal cap regions from analysis. **(C)** Cross section view of the hydrophobic cavity of RBGs 3-4, in space filling representation, showing density corresponding to lipids in the cryo-EM maps. Local resolution of lipids density is shown in the lower panel. **(D)** Cross section view of the bridge domain, highlighting the presence of lipids within the channel spanning from RBG3 to RBG10. **(E)** Representative cryo-EM maps of the VPS13A/CaM-XKR1 complex obtained from 3D classification, illustrating conformational changes in the VAB domain. The VAB adopts a continuum of conformations; the state shown in dark blue is more compatible with lipid transfer. **(F)** Representative frames from 3D variability analysis illustrating the coordinated movement of the VAB and C-terminal cap of VPS13A. Note the opening of the lipid transfer groove as the VAB adopts the more lipid-transfer compatible conformation.

The lipid transfer groove of VPS13s can accommodate tens of glycerophospholipids (Srinivasan et al., 2024). Based on previous mass spectrometry analyses of the lipids that co-purify with fungal Vps13 or human ATG2A (Kumar et al., 2018; Valverde et al., 2019), the bound lipids are glycerophospholipids, with varying acyl chain and head group compositions. The reconstruction represents an average of ∼0.4 million VPS13A/CaM-XKR1 complexes, so that the density for lipids in the transfer groove is also averaged. Nevertheless, we observe density interpretable as lipid in the best resolved parts of the bridge domain in RBGs 3-10 (Fig 2C-D). In particular, we can resolve rows of fatty acyl moieties along the hydrophobic walls of the taco-shell, indicating that there are distinct binding sites for fatty acyl chains within the bridge domain. The corresponding headgroups, which are expected to be solvent exposed at the top of the taco shell, as well as lipid tails in the middle of the transfer channel away from the walls, are not resolved in the maps.

The “cap” regions at either end of the bridge domain (Fig. 2A) are poorly defined in the maps of the VPS13A/CaM-XKR1 complex, reflecting high mobility, and we cannot confidently model them (Fig. 1D). The VAB domain is also mobile, as reflected by the low resolution in corresponding parts of the map (Fig. 1D). The particles used for reconstruction can be separated into different 3D classes, showing the VAB domain in a continuum of conformations (Fig. 2E-F), and its position in the model represents and average of these conformations. In the model the VAB domain is positioned at side of the bridge domain, allowing the bridge domain to connect to the acceptor membrane, in a “transfer-permissive” conformation. Notably, though, in the reconstruction of another VPS13 alone (VPS13C) (Li et al., 2025), not in complex with a membrane receptor like XKR1, the VAB is in a “transfer-incompatible” conformation, arching over the bridge-domain to prevent its connection to the membrane and thus maintaining VPS13 in a lipid-transfer incompatible form. In contrast to the C-terminal cap in VPS13A, the cap in VPS13C is well defined in the 3D reconstruction, showing that linker segments between the cap’s beta-strands interact with the VAB to maintain it in the transfer-inactive conformation. In the VPS13C structure (and in AlphaFold predictions for VPS13A), two alpha helices in the cap (residues 2799-2810 and 2839-2860 in VPS13A) partially obstruct the lipid transfer channel. 3D variability analysis of the particles used in the reconstruction of the VPS13A/CaM-XKR1 complex reveals a coordinated movement of densities corresponding to the VAB and the cap in the transition between conformations (Fig. 2E-F). The rearrangements in the cap are consistent with a potential role in facilitating lipid exit from the bridge domain (Fig 2F). We expect that as VPS13s engage the acceptor membrane, their VAB adjusts from a lipid-transfer inactive conformation, where the C-terminal cap is ordered, to transfer-active conformation, in which the cap rearranges to engage with membrane for lipid delivery. Likewise, we expect that the N-terminal cap could undergo structural changes as it engages with the donor membrane. Given the transfer-active and -inactive conformations of the VAB observed in our structure here and the VPS13C structure (Li et al., 2025), respectively, we propose that VPS13 activity in cells could be regulated via the transition between conformations that differ in the position of the VAB vis-à-vis the bridge domain.

Finally, we observed density that could not be accounted for by VPS13A alone in maps for its N-terminal end. Based on studies with other VPS13 proteins, this density corresponds to CaM (Li et al., 2025), which co-purifies with VPS13’s, including VPS13A (Sup. Fig. S1). The CaM binding site on VPS13s is near the end of the bridge domain that interacts with the ER, raising the possibility that CaM may be an exogenous adaptor for ER association (Li et al., 2025).

### Architecture of XKR1

Experimental structures of XK-related proteins (XKR4, XKR8, XKR9) have been reported before (Chakraborty et al., 2025; Sakuragi et al., 2021; Straub et al., 2021). Consistent with these structures and AlphaFold predictions (Jumper et al., 2021), XKR1 comprises two modules, each consisting of four TM helices (TM1-TM4 in module 1; TM5-TM8 in module 2) and “inter-helices” (IH 1-2 in module 1; IH 3 in module 2), which dip into but do not extend the whole distance through the lipid bilayer, between the second and third TM helix of each module (Fig. 3A).

**Figure 3.**
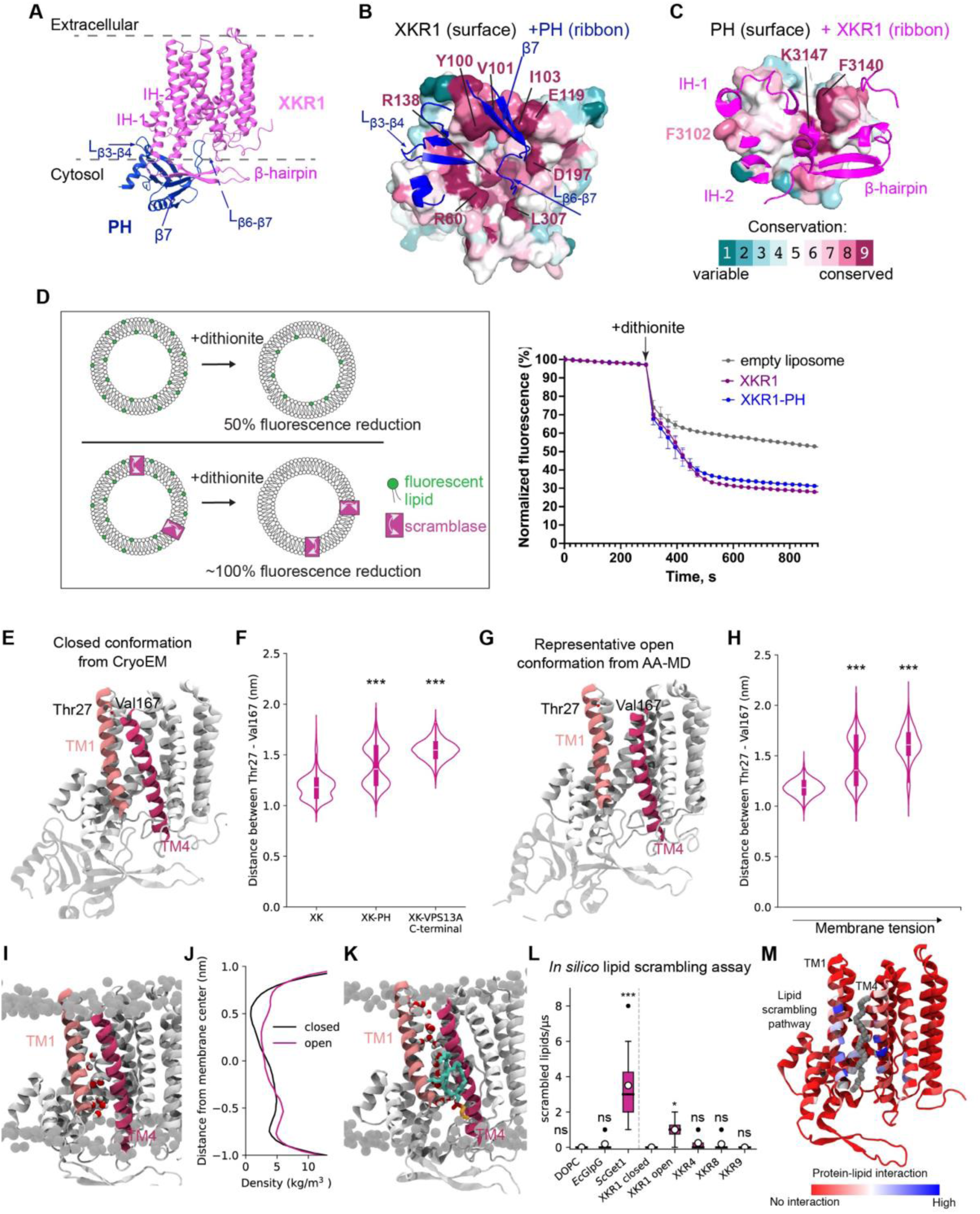
Lipid scrambling by XKR1-PH_(VPS13A)_ complex. **(A)** Model of the XKR1-PH_(VPS13A)_ complex as embedded in the plasma membrane (indicated). XKR1 is shown in pink and the PH domain in dark blue. Interaction region of both XKR1 and the domain are indicated. **(B)** Surface view of XKR1 viewed from the cytosol; PH domain segments at the interface with XKR1 are in ribbon representation. The β-hairpin and the cytosol accessible pocket are highly conserved. **(C)** Surface view of the PH domain as seen from XKR1. XKR1 segments at the interface with the PH domain are in ribbons representation. Key conserved residues at the XKR1:PH domain interface are labeled. Conservation scores of residues of the PH domain (B) and XKR1 (C) were analyzed via Consurf (https://consurf.tau.ac.il/consurf_index.php). **(D)** In vitro lipid scrambling assay, with schematic illustration on the left, shows that the XKR1-PH fusion protein demonstrated lipid scrambling activity comparable to XKR1 alone. **(E)** XKR1-PH_(VPS13A)_ complex in a closed conformation as based on the cryo-EM reconstruction. The TM helices 1 and 4 from XKR1 are shown in light pink and pink ribbons, respectively. The rest of the protein and the PH domain are shown in gray. **(F)** Distance between the Cα atoms from the residues Thr27 and Val167, located at the extracellular side of the TM1 and TM4, respectively, for unbiased systems. One way ANOVA with Tukey’s multiple comparison test; *** : p=0.001. **(G)** Snapshot of the XKR1-PH_(VPS13A)_ complex in an open conformation from AA-MD simulations, using the same color scheme as in panel (E). **(H)** Distance between the Cα atoms from the residues Thr27 and Val167 in systems with increased applied membrane tension, 0 mN/m, 10 mN/m, and 15 mN/m. One way ANOVA with Tukey’s multiple comparison test; *** : p=0.001. **(I-K)** Hydration analysis of the cavity/interface between TM1 and TM4. (I) Representative snapshot of the closed XKR1 conformation. (J) Water density profile along XKR1 for the closed (black) and open (pink) conformations. (K) Representative snapshot of the open XKR1 conformation showing a lipid located at the cytosolic entrance of the cavity/interface between TM1 and TM4. The lipid tails are shown as cyan sticks, phosphate atoms in red and choline atoms in orange. The protein is colored as in panel (E). **(L)** *In silico* lipid scrambling assay for experimentally available structures of XK-related proteins and the open conformation obtained from AA-MD simulations. One way ANOVA with Tukey’s multiple comparison test; *** : p≤0.001, * : p≤0.05, ns: p>0.05. **(M)** Key residues involved in lipid scrambling for the open conformation of XKR1. The averaged lipid scrambling pathway is shown as a continuous stalk of gray spheres. The red (no interaction) - blue (high interaction) color code indicates the interaction between each amino acid and the scrambled lipids.

XKR1 and all XK-related proteins feature a hydrophilic pocket facing the cytosolic surface of the membrane (Fig. 3B). In XKR1, the pocket is partially covered by a beta hairpin between IH2 and TM3 (Fig. 3A-C), which is not present in most other family members. Residues at the surface of the pocket, including in the beta hairpin in the case of XKR1, are highly conserved, indicating functional significance discussed further below.

### Interaction between VPS13A and XKR1

On its cytosolic side, XKR1 binds VPS13A exclusively via the conserved pocket (including the beta hairpin) that interacts with the PH domain of VPS13A through an extensive occluded surface area (∼2500 Å^2^) (Fig. 3B). PH domain residues at the interface are also conserved. The PH domain may be described as a sandwich of two antiparallel beta sheets, comprising strands 1-4 and 5-7, followed by a helix. Strand 7 of the PH domain associates with the XKR1 beta hairpin, extending the PH domain’s C-terminal beta-sheet. This interaction positions the PH domain directly over XKR1’s conserved pocket and was previously shown to be essential for VPS13A targeting to the plasma membrane (Guillen-Samander et al., 2022; Park et al., 2022). Further, loop PH-L _β3-β4_ of the PH domain is in the micelle, inserted between IH1 and IH2 of XKR1 adjacent to the conserved surface pocket, via a phenylalanine residue at its tip, F3102 (Fig. 3 A-C). F3102 is sandwiched between a leucine (L68) and a phenylalanine (F81) of XKR1 IH1 and IH2, respectively. A mutant construct, in which the sequences in PH-domain beta strands 3 and 4 and PH-L _β3-β4_ were replaced with the corresponding sequences from VPS13C, interfere with the PH domain’s targeting to the plasma membrane, consistent with functional importance (Guillen-Samander et al., 2022). And finally, intriguingly, the loop between strands 6 and 7 of the PH domain (PH-L _β6-β7_) extends into XKR1’s conserved surface pocket (Fig. 3A-C).

In structures of full-length XKR8 and −9 (but not XKR4) peptides from their C-terminus occupy this pocket (Chakraborty et al., 2025; Sakuragi et al., 2024; Sakuragi et al., 2021; Straub et al., 2021). Different from XKR1, XKR8 and XKR9 in cells are activated when caspases cleave off their C-termini to vacate the pocket, and, as such, the presence of the peptide in the pocket was speculated to be inhibitory (Sakuragi et al., 2024; Sakuragi et al., 2021; Straub et al., 2021), although such inhibition has never been demonstrated in a reconstituted system.

To test the PH domain’s effect on XKR1 scrambling, we carried out well established *in vitro* scrambling assays (Ploier and Menon, 2016) with FLAG-tagged XKR1 alone versus a FLAG-tagged XKR1-PH fusion construct, wherein the PH domain of VPS13A was fused to the XKR1 C-terminus. In these assays, candidate scramblases are reconstituted into liposomes containing trace amounts of NBD-labeled lipids, then a reducing agent (dithionite) is added. In liposomes lacking an active scramblase, dithionite reduces only NBD-lipids in the cytosol accessible leaflet of the liposome membrane, leading to a 50% reduction in NBD fluorescence. The reduction is larger when a scramblase is present to equilibrate NBD-lipids between the inner and outer leaflets, so that they are all dithionite accessible (Fig. 3D). As shown before (Adlakha et al., 2022), XKR1 by itself scrambles. Moreover, the XKR1-PH construct also scrambles, indicating that the binding of the PH domain and insertion of PH-L_β6-β7_ into XKR1’s conserved cytosolic pocket do not abrogate XKR1 scrambling activity (Fig. 3D). XKR1 therefore is active as a scramblase in the complex with VPS13A.

We propose that in XKR1 as well as in other XK-related proteins, the conserved surface pocket represents a binding site for protein partners, like VPS13A for XKR1. While activation of XKR-8 and −9 *in cellulo* correlates with removal of their C-terminal peptides from the conserved surface pocket, the function of the C-termini may not be inhibitory. For example, their removal in XKR8 and XKR9 might contribute to the recruitment of protein factors that collaborate with the XK-related proteins in their physiological function of apoptosis.

### XKR1 scrambling mechanism

The best characterized scramblases, such as those in the TMEM16 family, feature a hydrophilic groove facing the greasy membrane core that allows lipid head groups to slide through the membrane, like a credit-card through a reader (Pomorski and Menon, 2006; Sebinelli et al., 2024). We did not observe such a groove in the model of XKR1 derived from the Cryo-EM reconstruction, nor has such a groove been identified in the experimentally derived structures of other XK-related proteins (Chakraborty et al., 2025; Sakuragi et al., 2021; Straub et al., 2021).

As a first step to characterize the mechanism of lipid scrambling by XKR1, we performed all-atom molecular dynamics (AA-MD) simulations of the cryo-EM-derived XKR1 structure embedded in a lipid bilayer, either as a monomer, in the presence of the PH domain of VPS13A, or in the presence of the entire C-terminal region of VPS13A (see Sup. Table S3 for a detailed description of the simulated systems), including the PH domain. We observed that when bound to the PH domain of VPS13A, XKR1 spontaneously undergoes a conformational change with respect to the cryo-EM structure, characterized by a marked opening of the distance between TM1 and TM4 (Fig. 3E-G and Sup. Fig. S4). A similar transition is observed when the membrane in which XKR1 is embedded is subjected to increasing membrane tension (Fig. 3H). The opening between TM1 and TM4 is associated with an increased water permeation, and the binding of one lipid at the cytosolic side of the interface between TM1 and TM4, likely priming the phospholipid for exposure to the extracellular leaflet (Fig. 3I, K). However, no complete lipid translocations were observed in any of the AA-MD simulations, consistent with previous AA-MD simulations of XKR4 (Chakraborty et al., 2025). We will refer to the XKR1 conformation observed in the cryo-EM reconstruction as “closed” and the alternative conformation from the AA-MD simulations as “open”.

After conversion of these two conformations to coarse grain (CG) resolution to assess their lipid scrambling propensity, CG-MD simulations suggest that while the “closed” structure is not able to scramble lipids, the “open” structure can do so (Fig. 3L), and that the lipid scrambling pathway proceeds through a cavity between TM1 and TM4 lined with polar residues (Fig. 3M) . Although this channel is closed in all XKR cryo-EM structures, mutational analysis of XKR8 and XKR4 suggests that residues in TM1, TM4, and TM3, which is at the back of the cavity in the proposed open conformation, play roles in scrambling (Chakraborty et al., 2025; Sakuragi et al., 2021).

Overall, our simulations indicate that while the structure of XKR1 emerging from the cryo-EM analysis represents a scrambling-inactive state, its activation when membrane-embedded is likely to involve opening of a polar groove at the extracellular side of the interface between TM1 and TM4 helices, and this is likely to occur spontaneously when reconstituted in liposomes.

### VPS13A-XKR1 complex thins and deforms membranes

Lipid extraction from donor membranes and insertion into acceptor membranes pose energetic barriers that must somehow be overcome for efficient BLTP-mediated bulk lipid transfer. Therefore, we next examined the interaction of the VPS13A-XKR1 complex with membranes, to investigate whether the cryo-EM structure is compatible with lipid delivery and to identify factors that might facilitate lipid transport.

To investigate the interaction between the VPS13A-XKR1 complex and lipid membranes with high spatial resolution, we turned to CG-MD simulations of the entire complex embedded into a plasma-membrane like model lipid bilayer (see Sup. Table S2 for details on lipid composition). A challenge arising from the cryo-EM reconstruction is that some of the elements in VPS13A’s C-terminus that might mediate membrane association are disordered or highly mobile and could not be determined from the cryo-EM map. This is because the VPS13A sample in the cryo-EM reconstruction was not associated with membranes. Of particular interest among the disordered regions are the amphipathic helices of the ATG2_C motif, as they are known to play important roles in the localization of both VPS13 and ATG2 family BLTPs (Kumar et al., 2018; Wang et al., 2025b), and because amphipathic helices have the ability to bind and remodel membranes (Gimenez-Andres et al., 2018), which may play a role on lipid exchange.

After modelling the disordered regions of ATG2_C back into the VPS13A structure (see Methods), our simulations indicate that the architecture of the VPS13A-XKR1 complex is indeed compatible with direct interactions between VPS13A’s bridge domain and the cytosolic leaflet of the plasma membrane (Fig. 4A, B). Specifically, while anchored to XKR1 via its PH domain, VPS13A binds to the membrane via its ATG2_C motif, which directly embeds into the bilayer with its amphipathic helices (Fig. 4B). Deletion of the ATG2_C motif (but not of the PH domain) from full length VPS13A, disrupts this interaction, promoting dissociation of the VPS13A lipid-transport bridge from the bilayer (Fig. 4B). To validate this observation, we next tested the role of VPS13A’s ATG2_C motif in protein-membrane interactions, by performing liposome flotation assays. These show that the ATG2_C motif supports membrane association of VPS13A (Fig. 4C).

**Figure 4.**
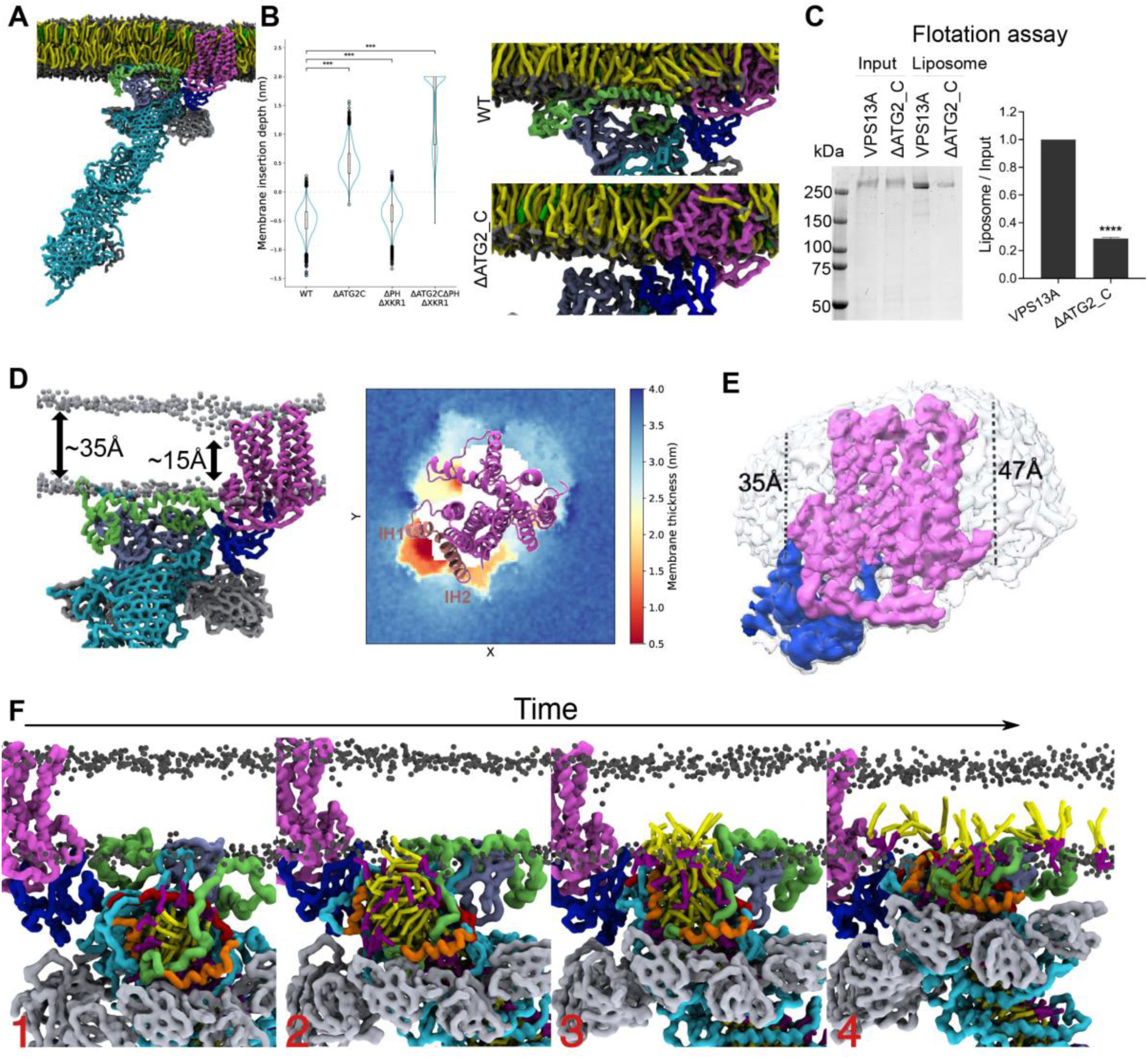
VPS13A-XKR1 complex interactions with membranes and supports bulk lipid delivery. **(A)** Snapshot of the CG model of VPS13A/CaM-XKR1 complex embedded in a plasma membrane (PM)-like membrane, the color scheme of the different domains and monomers matches that used in Figure 1. **(B)** Insertion depth (in nm) of wild-type VPS13A and mutants into the membrane, computed as the difference between protein’s *z* coordinate and the local *z* coordinate of the bilayer headgroups. Representative snapshots (right panel) show the membrane interface between of wild-type VPS13A and the ΔATG2_C mutant. Statistical analysis was performed using ordinary one-way ANOVA and Welch’s pairwise comparisons with Bonferroni correction (*** P<0.001, 2 replicates). **(C)** Membrane association of full-length VPS13A and ΔATG2_C mutant analyzed by biochemical flotation assay. A representative SDS-PAGE gel shows reduced flotation of ΔATG2_C mutant with liposomes compared to wild-type VPS13A. Quantification from three independent experiments is shown in the right panel. Data are presented as mean ± SD. Statistical analysis was performed using an unpaired Welch’s t-test (**** P <0.0001). **(D)** Snapshot and membrane thickness *xy* map from CG-MD simulations showing local membrane thinning around helices IH1 and IH2 of XKR1. **(E)** Cryo-EM density of XKR1-PH complex, with XKR1 embedded in GDN detergent micelles visualized as a transparent white density. The micelle thickness was measured at two opposing sides surrounding XKR1. **(F)** Time-evolved snapshots (panels 1-4) of a CG-MD simulation with the VPS13A hydrophobic groove pre-loaded with 93 lipids, showing bulk lipid delivery. Delivered lipids are colored in yellow, with headgroups in purple; phosphate groups of the membrane lipids are displayed as black spheres.

Further, our CG-MD simulations show that the presence of XKR1 causes significant remodeling of the surrounding membrane environment. First, consistent with observations for other scramblases, XKR1 promotes local membrane thinning (∼15 Å) (close to helices IH1 and IH2 (Fig. 4D, Sup. Fig. S5)(Chakraborty et al., 2025; Sebinelli et al., 2024). Such distortion parallels observations from the cryo-EM reconstruction, where the detergent micelle surrounding XKR1 is asymmetric, with a ∼25% thinning around XKR1 IH1 and IH2, close to where the VPS13A bridge-domain is expected to interact with the membrane (Fig. 4E). Moreover, the presence of XKR1 induces non-negligible membrane curvature in its surroundings (Sup. Fig. S6). This curvature is further enhanced by the ATG2_C motif of VPS13A (Sup. Fig. S6), indicating cooperation between the various components of the complex towards membrane remodeling.

### Direct lipid transfer from the VPS13A bridge domain to the proximal membrane

To investigate the molecular details of lipid transfer from VPS13A’s bridge domain to the membrane, we next performed unbiased CG-MD simulations of the complex in the presence of different amount of lipids (54, 73 and 93) inside the hydrophobic cavity of VPS13A, as a proxy to model lipid synthesis from the ER. When the lipid cavity was not fully occupied (54 and 73 lipids, respectively) no spontaneous delivery of lipids to the proximal membrane was observed, even if the protein was fully bound to the lipid bilayer via its ATG2_C motif (as in Fig. 4A,B). Estimation of the energy barrier required for such transfer by means of a potential of mean force (PMF), indicates an upper limit of around 10 kcal/mol for the lipid transfer barrier (Sup. Fig. S7). In contrast, we observed extensive lipid transfer (up to 27 lipids) in multiple unbiased simulations in the presence of excess bound lipids (Fig. 4F, Sup. Fig. S8, Movies S1 and S2). In detail, the excess lipids start gathering near the C-terminal cap of the protein, with the headgroups protruding out of the hydrophobic cavity and facing the membrane (Fig. 4F, panel 1). Simultaneously, consistent with the 3D variability analysis, helices present in the cap (residues 2799-2810, 2839-2860) are mobile, opening the lipid transfer groove toward the membrane (Fig. 4F, panel 2). Eventually, the gathered lipids make contact with the membrane, and they are delivered in bulk (Fig. 4F, panels 2, 3). In multiple replicas, this process occurs concomitantly with significant conformational transitions of the cap helices, which start interacting with the membrane as well, and of the ATG2_C motif (Fig. 4F, panels 3, 4). Increasing membrane tension in the acceptor bilayer accelerates the delivery process, by promoting closer interaction between the bridge domain’s C-terminal cap and the bilayer (Sup. Fig. S9) and likely by membrane stabilization due to lipid delivery into the tense membrane.

That XKR1 more readily and stably adopts its “open”, lipid-scrambling competent, conformation in the presence of high membrane tension (Fig. 3H , Sup. Fig. S4) suggests that lipid delivery by VPS13A and scrambling activity by XKR1 could be coupled by physical or chemical modifications of bilayer properties, even in the absence of a direct physical connection between the bridge domain and XKR1.

## Discussion

Together, our experimental structure of the VPS13A/CaM-XKR1 complex and MD analysis suggest that lipid delivery is activated when VPS13A interfaces with its binding partner XKR1 in the acceptor membrane, and then lipids are delivered directly from the bridge-domain of VPS13A to the cytosolic leaflet of the plasma membrane. The interaction between VPS13A’s PH domain and XKR1, conformational changes in VPS13A’s C-terminal “cap” and its VAB domain, and membrane insertion of the amphipathic helices of VPS13A’s ATG2_C motif are required to juxtapose the end of the bridge-domain to the proximal membrane, so that lipids can move from the hydrophobic environment within the bridge to the membrane without desolvation of their acyl chains. Our studies suggest that the VPS13A-XKR1 complex introduces significant local membrane deformations in the acceptor membrane where lipid transfer from the VPS13A bridge is expected to take place. These deformations impact lipid packing and may lower the energy barrier for lipid insertion into the acceptor membrane (Álvarez et al., 2025).

Importantly, we were able to observe robust lipid transfer in silico when lipids were present in excess inside the bridge domain, a condition that mimics lipid synthesis at the ER. Further, lipid release was accelerated by an increase in membrane tension at the acceptor membrane. These results support current thinking that directional BLTP-mediated lipid flow could be driven by differences in membrane tension between donor and acceptor membranes (Reinisch et al., 2025; Zhang and Lin, 2024). The ER, the donor membrane where most structural lipids are synthesized, has low membrane tension, whereas membrane tension in other cellular organelles is higher (Wang et al., 2025a). How the differences in membrane tension arise is not currently well understood and is a topic attracting significant attention (Barahtjan et al., 2025).

VPS13A delivers its lipids to the cytosolic leaflet of the membrane close to XKR1, which can then equilibrate newly delivered lipids between leaflets of the bilayer. As assessed *in silico*, the XKR1 conformation observed in the cryo-EM reconstruction does not support lipid scrambling, nor do experimental structures for other XK family members (Chakraborty et al., 2025; Li et al., 2024). AA-MD simulations for XKR1 suggest, however, that another conformation exists, where TM1 and TM4 have moved apart to form a polar channel through which lipids can slide between bilayer leaflets. Both XKR1 and a XKR1-PH fusion construct scramble lipids when embedded in liposomes in vitro, supporting that XKR1 may adopt a scrambling active conformation in membranes, versus in a detergent micelle as in the cryo-EM reconstruction. This observation parallels observations for TMEM16 scramblases at the plasma membrane, that also undergo opening/closing of their hydrophilic grooves to regulate lipid scrambling activity (Feng et al., 2024; Khelashvili et al., 2022; Stephens et al., 2025). Mutational analyses of XKR4 and XKR8 are consistent with the proposal that a groove between TM1 and TM4 opens to enable scrambling, but whether the remaining members of the XK-related family (XKR2, XKR3, XKR5-7) act by the same mechanism remains to be investigated.

In the VPS13A-XKR1 complex under study here, its PH domain plays a primary role in targeting VPS13A to XKR1 at the plasma membrane, where it coordinates with the ATG2_C motif in firmly anchoring the bridge domain to the membrane. We cannot exclude that the VAB and WWE domains may assist in membrane binding via additional interactions. Moreover, VPS13A also functions at other cellular locations (Kumar et al., 2018) together with receptors other than XKR1, where the VAB or WWE domains or some combination of the PH, VAB, or WWE domain could mediate organelle attachment. We would envision that the VAB or WWE domains could have the same function as the PH domain in positioning the bridge-domain with respect to the acceptor membrane. The VPS13 proteins, including VPS13B, -C, -D in humans as well as those in other organisms, share a similar architecture, with shared built-in adaptor modules (ATG2_C, PH, VAB; the WWE domain is present only in VPS13A and C). Thus, the structure of the VPS13A/CaM-XKR1 complex we describe here is applicable to understand lipid delivery by all VPS13 proteins.

The ATG2 proteins, which are related to the VPS13s, lack most of the built-in adaptor domains of the VPS13 family, and instead they rely on exogenous partner proteins for their localization. They are streamlined versions of VPS13, pared down to the bridge-domain and the ATG2_C amphipathic helices. ATG2 cooperates with the scramblase ATG9 during autophagy and can bind to it directly (Ghanbarpour et al., 2021; van Vliet et al., 2022). Based on our current study of VPS13A lipid delivery, we propose that lipids are transferred from the bridge-domain of ATG2 directly to membrane, where the ATG2_C motif and the ATG2-ATG9 interaction serve to appose the bridge domain close to the acceptor membrane while also localizing the BLTP near a scramblase. A recent molecular dynamics study supports that ATG2 could deliver lipid directly to the acceptor membrane and describes a role for the ATG2_C amphipathic helices in membrane deformation during this process (Sakai et al., 2025). Worth noting, in cells during autophagy, ATG9 localizes to the rim of the nascent autophagosome (Melia et al., 2020), which is highly curved.

In contrast to our model, earlier studies of the ATG2-ATG9 complex at low resolution had led to speculation that ATG2 delivers lipids to the scramblase rather than directly to membrane and that the scramblase then integrates the lipids into the membrane (van Vliet et al., 2022; Wang et al., 2024). Another interpretation of the earlier data is consistent with the model put forth here: that ATG2 is tethered to ATG9 to bring it to the membrane in the proximity of scrambling activity, even as lipids are transferred from the bridge domain directly to the membrane. Thus, our study significantly advances our understanding of lipid delivery by BLTPs in the ATG2 family. Other less studied BLTP families, whose members also lack the ATG2_C amphipathic helices, could also deliver lipids directly to the acceptor membrane, although the features responsible for membrane binding and deformation remain to be identified.

A recent atomic resolution study of a BLTP1-containing complex that forms at the donor membrane (ER) showed that lipids enter the BLTP1 bridge-domain via an integral membrane protein vestibule, consisting of five TM helices (Kang et al., 2025). Our study here complements the BLTP1 study insofar as we describe mechanisms for lipid transfer out of the bridge domain and back into the membrane. Arguably unexpectedly, the lipid transfer process at the acceptor membrane, where at least for VPS13s lipids are transferred directly to the membrane, does not mirror the extraction process at the donor membrane observed for BLTP1. We look forward to further studies of BLTP proteins and processes at both ends that should help understand how general each of these strategies are.

## Data & Code Availability

Cryo-EM maps deposited in the EMDB (accession code EMD-72912, EMD-72909, EMD-72913) and coordinates derived from each of the maps deposited in the PDB (PDB ID 9YG4, 9YFW, 9YG5) are hold for release.

## Acknowledgments

We thank H. Zhou and S. Hamill for technical assistance. This work was funded by the NIH (R35GM131715 to KMR and R01DA018343 to PDC), by the Swiss National Science Foundation (grant <u>CR00I5-236020</u> to SV), and by the European Research Council under the European Union’s Horizon 2020 research and innovation program (grant agreement no. 803952, to SV). This work was supported by grants from the Swiss National Supercomputing Centre under projects ID s1269, lp24 and lp69. DA acknowledges support from the Margarita Salas program 2021–2023 funded by Ministerio de Universidades (MU-21-UP2021-030-53773022).

## Author Contributions

KMR, BH, and SV conceived of and supervised this project. BH carried out the cryo-EM analysis of the VPS13A/CaM-XKR1 complex and biochemical experiments, CRR investigated XKR1 scrambling activity *in silico*; VG determined the effect of the complex on membranes *in silico*; DA performed *in silico* investigation of lipid transfer. DL modeled the VPS13A N-terminus and its interaction with CaM and shared his coordinates for the lipid transfer inactive VPS13C complex before publication. XW and PDC were invaluable in discussions. KMR and SV wrote the manuscript, with input from all authors.

## Methods

### Protein expression and purification

To purify VPS13A protein, the codon-optimized VPS13A gene was cloned into a pCAG expression vector containing a C-terminal 3xFLAG tag. For the VPS13A ΔATG2_C mutant construct, the region encoding the ATG_2C motif (amino acid residues 2866-3029) was deleted from the full-length VPS13A sequence. The resulting truncated construct was cloned into the same pCAG vector backbone. The appropriate plasmid construct was transfected into Expi293 cells with ExpiFectamine (Gibco), following manufacturer’s instruction. 48h post transfection, cells were harvested, resuspended in lysis buffer A (50mM HEPES, pH7.4, 300mM NaCl, 1mM TCEP, 10% glycerol, 0.1% Triton X-100) supplemented with protease inhibitor cocktail (Roche). The suspension was incubated on a rotary shaker at 4°C for 15min, followed by mechanical lysis using a Dounce homogenizer (15-20 passes). The crude lysate was centrifuged at 39,191g for 30min at 4°C to remove cell debris. The supernatant containing the extracted protein was incubated with pre-washed anti-FLAG resin (#A2220, Millipore) for 2h at 4°C. The resin was then washed three times with buffer B (50mM HEPES pH7.4, 300mM NaCl, 1mM TCEP, 10% glycerol), followed by incubation in buffer B supplemented with 2mM MgCl_2_ and 1mM ATP at 4°C overnight to remove chaperone. After three additional washes with buffer B, the bound protein was eluted from the resin using 0.25 mg/mL 3xFLAG peptide (APExBIO) in buffer B supplemented with protease inhibitors. The eluted protein was concentrated through a 100 kDa molecular weight cutoff Amicon centrifugal filter and analyzed by SDS-PAGE stained with Coomassie blue. Calmodulin (CaM) co-purified with VPS13A, resulting in a native VPS13A/CaM complex.

The expression and purification of 3XFLAG-XKR1 was performed as previously described (Adlakha et al., 2022) with slight modifications. . To make a 3XFLAG-XKR1-PH fusion construct, the region encoding the VPS13A PH domain (amino acids residues 3031-3174) was fused to the C-terminus of XKR1. The resulting construct was cloned into the same pCMV vector with a N-terminal 3xFLAG. Expi293 cells were transfected with plasmids encoding XKR1 or the XKR1-PH fusion construct, and were harvested 48h post transfection. Cell pellets were resuspended in lysis buffer C (50mM HEPES, pH8, 200mM NaCl, 1mM TCEP, 10% glycerol) supplemented with protease inhibitors and lysed using a Dounce homogenizer. To extract XKR1 from membrane, GDN (Anatrace) was added to the lysate to a final concentration of 1.5%, followed by incubation on a rotary shaker at 4°C for 1.5 hours. The lysate was then clarified by centrifugation, and the supernatant was incubated with anti-FLAG resin pre-washed with buffer C containing 0.02% GDN. The remaining purification steps followed the protocol used for VPS13A, with 0.02% GDN included throughout. The final XKR1 protein was concentrated using a 10kDa molecular weight cutoff Amicon centrifugal filter and quantified by Coomassie blue staining using BSA standards.

To allow formation of VPS13A/CaM-XKR1 complex, an excess amount of XKR1 was incubated with purified VPS13A/CaM for 30min at room temperature. The mixture was then loaded onto a Superose 6 Increase 10/300 GL size-exclusion chromatography (SEC) column (GE Healthcare) pre-equilibrated with sample buffer (50mM HEPES, pH8, 200mM NaCl, 1mM TCEP, 0.02% GDN). Peak fractions corresponding to the VPS13A/CaM-XKR1 complex were pooled (Sup. Fig. S1A), concentrated using a 100 kDa Amicon centrifugal filter, analyzed by SDS-PAGE and used for cryo-EM grid preparation.

### Cryo-EM sample preparation and data collection

Freshly purified VPS13A/CaM-XKR1 complex was concentrated to 2.5-3.5mg/mL and supplemented with 0.1% Amphipol A8-35 (Anatrace) prior to grid preparation. A volume of 3.5µL of the complex was applied to a glow-discharged UltraAuFoil R1.2/1.3 300 mesh gold grids (Quantifoil), then plunge-frozen in liquid ethane using an FEI Vitrobot mark IV (Thermo Scientifc) under 100% humidity at 8°C, with a blot force of 1 and blot time of 3s. Data collection was carried out on a 300-kV FEI Titan Krios G2 transmission electron microscope equipped with a K3 summit direct detection camera at the Yale CryoEM Resource. A total of 26434 micrographs were collected in super-resolution mode at a nominal magnification of 105k (0.832 Å/physical pixel), and each movie was recorded with a total exposure of 1.4s fractionated into 50 frames for a total dose of 50 e^−^/Å^2^ and with a defocus range of −0.8 to −2 µm.

### Image processing

Raw micrographs were imported into CryoSPARC (v4.6.2) for processing (Sup. Fig. S2). Beam-induced motion was corrected using the Patch Motion Correction algorithm with a Fourier cropping factor of 1/2, and defocus values were estimated using Patch CTF Estimation with default parameters. Micrographs with a CTF fit resolution worse than 6Å were excluded from downstream processing.

Initial 2D templates were derived from blob picking followed by two rounds of 2D classification. A subset of 1662 micrographs was then used for template picking and 669,210 particles of box size 672 pix with 4x Fourier cropping were extracted. After further 2D classification, 101,081 particles were selected to generate a primary 3D reference volume using *ab initio* reconstruction. To obtain an improved reference volume, a larger dataset of 3,814,669 particles from 13,751 micrographs were subjected to three rounds of heterogeneous refinement, using the initial reference and four additional “junk” volumes. This workflow yielded 287,147 unbinned particles, which produced a 3.85Å map via non-uniform (NU) refinement. The resulting volume was subsequently used for 2D classification and selection of 2D templates with a broader range of projected views.

Using the full dataset comprising 25,832 micrographs, particles obtained from template picking were subjected to 2D classification and two rounds of heterogeneous refinement, using the 3.85 Å reference volume and 4 “junk” volumes. Following refinement, approximately 1million unbinned particles were re-extracted and used for *ab initio* reconstruction. Using default *ab initio* settings with 4 classes, followed by heterogeneous refinement, 464,055 particles were selected for NU refinement, yielding a 3.41Å map focused on VPS13A/CaM complex. To resolve the VPS13A-XKR1 complex, the same particle set was subjected to *ab initio* reconstruction with the following settings: 4 classes, maximum resolution of 4 Å, initial resolution of 12 Å, and initial and final minibatch size of 500. After three rounds of heterogeneous refinement, a class of 385,639 particles centered on VPS13A-XKR1 were subjected to NU refinement, resulting a 3.46 Å map. Local refinements were subsequently performed to obtain focused maps of the VPS13A N-terminal region bound to CaM (VPS13A/Nt-CaM), the central bridge domain, and the C-terminal region of VPS13A in complex with XKR1 (VPS13A/Ct-XKR1).

To analyze conformational heterogeneity of the C-terminal region of VPS13A, a focused 3D classification was performed using a manually created mask around the C-terminus. 3D variability analysis (3DVA) was also conducted with the filter resolution set to 6 Å, using a “simple” output mode with 20 frames to visualize structural dynamics.

### Model refinement and validation

The initial model of VPS13A, XK-PH(VPS13A) complex, and VPS13A/Nt-CaM complex were predicted using AlphaFold 3 (https://alphafoldserver.com) (Abramson et al., 2024) then subjected to rigid body fitting in UCSF ChimeraX (Meng et al., 2023), followed by a deep-learning-based model rebuilt and fitting using DeepMainmast (Terashi et al., 2024), except that the VPS13A VAB domain (residues 1876-2504) was refined further due to low resolution. The models were then manually readjusted in Coot (Casanal et al., 2020) against corresponding focused maps derived from local refinement, followed by real space refinement in Phenix (Liebschner et al., 2019).

The following regions of VPS13A (1-33,50-53,128-139, 415-457, 609-625, 794-875, 898-902, 1015-1045, 1192-1201, 1342-1387, 1532-1549, 1671-1715, 1810-1827, 1867-2517, 2731-2764, 2784-2820,2832-3030, 3170-3174), XKR1 (267-269,388-444) and CaM (residues 1 and 149) were not modelled, as the corresponding densities were not visible or at low resolution in all the maps. Representative densities for different parts of the VPS13A/CaM-XKR1 complex are shown in Sup. Fig. S3.

Molecular graphics were prepared using UCSF ChimeraX (Goddard et al., 2018) and PyMOL (The PyMOL Molecular Graphics System, Version 3.0 Schrödinger LLC).

### Liposome preparation and flotation assay

To prepare liposomes mimicking the plasma membrane, a lipid mixture containing 30% DOPC, 27.5% DOPE, 12% DOPS, 30% Cholesterol, 0.5% NBD-PE (all in chloroform) was dried under a gentle N2 stream to a thin film, followed by vacuum desiccation overnight. The dried lipid film was rehydrated in buffer containing 50mM HEPES (pH7.4) and 200mM NaCl to a final lipid concentration of 5mM. The lipid suspension was incubated at 37°C for 1h and then subjected to 10 cycles of flash freeze-thaw. Crude liposomes were extruded 21 times through a 200 nm polycarbonate filter using a mini extruder (Avanti) to yield uniform vesicles.

To analyze liposome binding of VPS13A and the ΔATG2_C mutant, proteins were purified from Expi293 cells as described above. Seventy microliters of protein sample were incubated with 100µL 2.5mM liposome for 30min at room temperature. The mixture was adjusted to 30% OptiPrep by adding an equal volume of 60% OptiPrep (D1556, Sigma-Aldrich) and overlaid with 180µL 15% OptiPrep and 180µL reconstitution buffer (50mM HEPES, pH7.4 and 200mM NaCl) in a 0.8mL ultracentrifugation tube. After centrifugation (SW55Ti, 35,000 rpm, 1h, 16°C), the liposome-containing fraction at the 0%|15% interface was collected, and the liposome-bound proteins were analyzed by SDS-PAGE.

### Scrambling assay

Proteoliposomes, containing either XKR1 or the XKR1-PH fusion construct, were prepared as previously described (Li et al., 2024) with minor modifications. Fusing the PH domain to the XKR1- C-terminus rather than adding soluble VPS13A or the PH domain to XKR1-containing liposomes had two rationales: (i) since the XKR1 C-terminus is on the same side of the membrane as the regulatory pocket, this ensures that the PH domain can access the regulatory pocket irrespective of XKR1’s orientation after reconstitution into the liposome membrane and (ii) the fusion ensures that there are no XKR1s in a PH-domain-free form. Briefly, 180µL of 2.5mM liposomes (30% DOPC, 27.5% DOPE, 12% DOPS, 30% Cholesterol, 0.5% NBD-PE, as above) were mixed with 0.3% Triton X-100 for 30min at 37°C to partially solubilize the lipid bilayer. Subsequently, 20µL of concentrated protein was added and incubated for 1h at room temperature. Detergent was then removed by sequential addition of pre-washed Bio-Beads (Bio-Rad): 20mg for 1h at RT, followed by another 20mg for 1h at RT, and finally 40mg overnight at 4°C. Reconstituted proteoliposomes were then purified via OptiPrep flotation assay as described above.

To analyze lipid scrambling activity, 5µL proteoliposomes were added to 95µL of assay buffer (50mM HEPES, pH7.4, 200mM NaCl, 1mM TCEP) in a white 96-well plate (Corning). Each condition was measured in triplicate. NBD fluorescence was recorded using a plate reader (BioTek) with excitation/emission set to 460/538nm. After signal stablization, 5µL freshly prepared dithionite was added to a final concentration of 5 mM to initiate quenching of externally exposed NBD-lipids. Fluorescence was monitored for 10 min,followed by addition of 5µL of 10% Triton X-100 and 5µL dithionite to fully quench remaining fluorescence.

### System setup and simulation details

The starting structure of the VPS13A-calmodulin-XKR1 complex for both AA and CG MD simulations was taken from the cryo-EM refined structure. Missing loops in the structure were modelled using Modeller (Webb and Sali, 2021). The loop comprising residues 1671-1715 and the ATG2_C domain were aligned from the AlphaFold prediction (Abramson et al., 2024; Jumper et al., 2021), and the disordered ATG2_C domain was subsequently manually displaced from its initial position under the C-Cap of VPS13A. The complex was minimized in vacuum using the CHARMM36m force field (Huang et al., 2017) before running any AA or CG MD simulation. Both AA and CG MD simulations were carried out using the software GROMACS (https://doi.org/10.1016/j.softx.2015.06.001).

#### CG-MD Simulations

The atomistic structure of the protein complex was converted to a CG model using the Martinize2 script (https://doi.org/10.7554/eLife.90627.3). CG simulations were carried out using the force field (Souza et al., 2021) . An elastic network with a force constant of 700 kJ mol^-1^ nm^-2^, with an upper elastic bond cutoff of 0.9 nm, was used to restrain the secondary structure of the protein and the protein-protein interfaces. To reproduce the expected mobility of certain regions of the protein, the elastic network was removed *i)* from all loops, *ii)* between the ATG2_C domain and the rest of the protein, *iii)* between the two helices at the C-terminal region (residues 2790-2810, and 2839-2856) and the rest of the protein.

To fill the hydrophobic cavity of the protein with 54 lipids, the unbiased CG-MD protocol for iterative lipid binding from (Srinivasan et al., 2024) was followed, after aligning the published Vps13 structure with 49 lipids inside. To oversaturate the cavity with more lipids, we adapted a recently proposed *in silico* lipid synthesis protocol (Nieto et al., 2023) which consists of the following iterative steps: 1. Duplicating one of the lipids inside the cavity, 2. Slow-growth simulation of the duplicated lipid to avoid clashes, and 3. Standard equilibration + production for 50 ns.

The orientation in the membrane for the proteins was computed using PPM (Lomize et al., 2012), and the CG structure was embedded in a lipid bilayer using the Insane script (Wassenaar et al., 2015). Description of lipid composition, system size and simulation time are collected in Sup. Table S2. The systems were solvated and ionized with 150 mM NaCl; additional Na^+^/ or Cl^-^ ions were added to neutralize the charge of each system.

The standard minimization and equilibration protocol from CHARMM-GUI was used for each system, which consists of an initial minimization using the steepest descent algorithm followed by six NPT equilibrations with positional restraints on both the protein backbone beads and the lipid headgroups, which were gradually removed. The time step was initially fixed at 1 fs and gradually increased until 20 fs. The equilibration runs used the v-rescale thermostat and C-rescale barostat to maintain the temperature at 310 K and the pressure at 1 bar. For production, two independent replicas were run for 5 microseconds for each system using a time step of 20 fs. The v-rescale thermostat, with separate coupling for the protein, the lipids, and the ionized solvent, and a coupling time constant of 1.0 ps, was used to maintain the temperature during production. The semi-isotropic Parrinello-Rahman barostat, with a compressibility of 3*10^-4^ bar, a coupling time constant of 12 ps, and a 10-step frequency for the coupling was used to control the pressure at 1.0 bar during production runs. Electrostatic interactions were computed using the reaction-field electrostatics approach with a cut-off of 1.1 nm. Van der Waals interactions were computed using a cut-off 1.1 nm along with a potential-shift-verlet scheme. The neighbor list was updated every 20 steps using the Verlet list scheme.

#### AA-MD Simulations

The CHARMM-GUI Membrane Builder (Jo et al., 2009) was used to prepare the AA systems, with the membrane compositions listed in Sup. Table S3. Each system was solvated with CHARMM TIP3P water and ionized with 0.15 M of NaCl and neutralizing Na^+^ or Cl^-^ ions. The CHARMM36m force field was used (Huang et al., 2017). The CHARMM-GUI protocol for minimization and equilibration was followed for each system, which consists in an energy minimization using the steepest descent algorithm followed by two NVT equilibrations and four NPT equilibrations with positional restraints on both the protein backbone and the phosphate atom of each lipids that are increasingly removed. The equilibration runs used the v-rescale thermostat and the C-rescale semi-isotropic barostat to maintain the temperature at 310 K and the pressure at 1 bar. The v-rescale thermostat, with separate coupling for the protein, the membrane, and the ionized solvent, and a coupling time constant of 1.0 ps, was used to keep the system at 310 K during the production runs. The pressure of 1.0 bar in the productions runs was kept with the semi-isotropic C-rescale barostat, with a compressibility of 4.5·10^-5^ bar, and a coupling time constant of 5.0 ps. For the electrostatics, the Particle Mesh Ewald method was used to compute the electrostatic interactions, with a Fourier spacing of 0.12 nm and a cutoff of 1.2 nm. Van der Waals interactions were switched to zero over the 1-1.2 nm range. The LINCS algorithm was used to constrain the bonds involving hydrogen atoms (Hess et al., 1997). Frames were written every 100 ps, with a time step of 2 fs.

For the AA-MD and CG-MD simulations with tension on the membrane, NPT simulations with the C-rescale barostat and *surface-tension* as pressure coupling type were used, taking 1 bar on the z direction, and either 400 bar nm (20 mN m^-1^), 300 bar nm (15 mN m^-1^), or 200 bar nm (10 mN m^-1^) on the x/y dimensions of the box.

#### Umbrella-sampling simulations

The reaction coordinate for the umbrella sampling simulations was taken as the distance of the center-of-mass (COM) of the delivered lipid along the bilayer normal (z) to the z coordinate of the membrane center, which was calculated as the COM of all lipids contained within a cylinder of 2.0 nm radius around the COM of the delivered lipid. The windows were spaced at 0.1 nm, with a distance range between 1.2 and 4.5 nm. Each window was simulated for 505 ns, using a harmonic force constant of 500 kJ mol^-1^ nm^-2^. The first 205 ns were not considered, and the PMF was constructed using WHAM (Hub et al., 2010) as implemented by GROMACS. Convergence was tested by computing the PMF in 100 ns blocks after discarding the first 5 ns. The windows were selected from an initial pulling simulation along the reaction coordinate, using a force constant of 1000 kJ mol^-1^ nm^-2^ and a pulling rate of 2·10^-5^ nm ns^-1^.

### Analysis

The minimum distance between the membrane and the lipids inside the cavity was computed using the gmx mindist tool. The density of the different groups along Z was calculated using the gmx density tool, normalizing all densities so they integrate to 1. Visualization of the trajectories and rendering was done using VMD (Humphrey et al., 1996).

To count the number of lipids delivered, the distance along z between a terminal lipid tail bead of the POPC lipids (C4A) to the membrane center was computed for each frame. Noise was reduced by applying a block averaging technique every 10 ns. A lipid is considered as delivered when the averaged distance is less than 2.3 nm.

The density of the phosphate groups of membrane and RBG lipids as well as the C-cap of VPS13A along z was computed using the gmx density tool, only considering the simulation time between 1 μs and the time at which bulk lipid delivery happens. All mass densities where normalized so they integrate to 1.

The membrane insertion depth of the protein was calculated by measuring all pairwise distances between VPS13A beads and lipid glycerol beads. Then, for each time frame, and only considering the glycerol beads within 2.0 nm of the protein, *i)* the maximum (in absolute value) z-position of the protein beads and *ii)* the mean z-coordinate of the selected glycerol lipid beads were stored and subtracted to get the maximum membrane insertion depth of the protein per frame.

Membrane thinning was analyzed using python scripts with libraries MDAnalysis, Numpy and SciPy. MD trajectories were initially fitted to the backbone beads of the transmembrane part of the systems on x and y directions. Next, for each frame the z-coordinates of the phosphate beads were stored. The membrane thickness was obtained by subtracting the z-coordinates of the inner leaflet from the z-coordinates of the outer leaflet. The z-coordinates of the phosphate beads corresponding to the cytosolic leaflet were also used to compute membrane curvature, by subtracting the z coordinate of each bead and the average z coordinate of the entire leaflet for each frame. The results of both membrane thickness and curvature were averaged using a grid along x and y with a size of 0.1 nm.

The opening of the interface between the TM H1 and TM H4 from XKR1 were traced by measuring the distance between the carbon-α from the amino acids Thr27 and Val167. Distances were computed with the gmx distance tool. Data points were stored every 0.5 ns.

Lipid scrambling was quantified by tracking the tilt angles (θ) of each lipid with respect to the membrane plane normal throughout the simulation, following the same protocol as in (Li et al., 2024). Here, a flip (movement from the upper to the lower leaflet) or flop (movement from the lower to the upper leaflet) event was considered when the lipid exhibited a change to tilt angles θ ≥ 130° or θ ≤ 50°, respectively. The angles were computed every nanosecond with the gmx gangle tool, and events were reported as events/μs, omitting the first 2μs for equilibration. Data for the negative (pure DOPC membrane and EcGlpG) and positive (ScGet1) controls can be found in https://doi.org/10.5281/zenodo.10475371.

## Supplementary data

**Figure S1.**
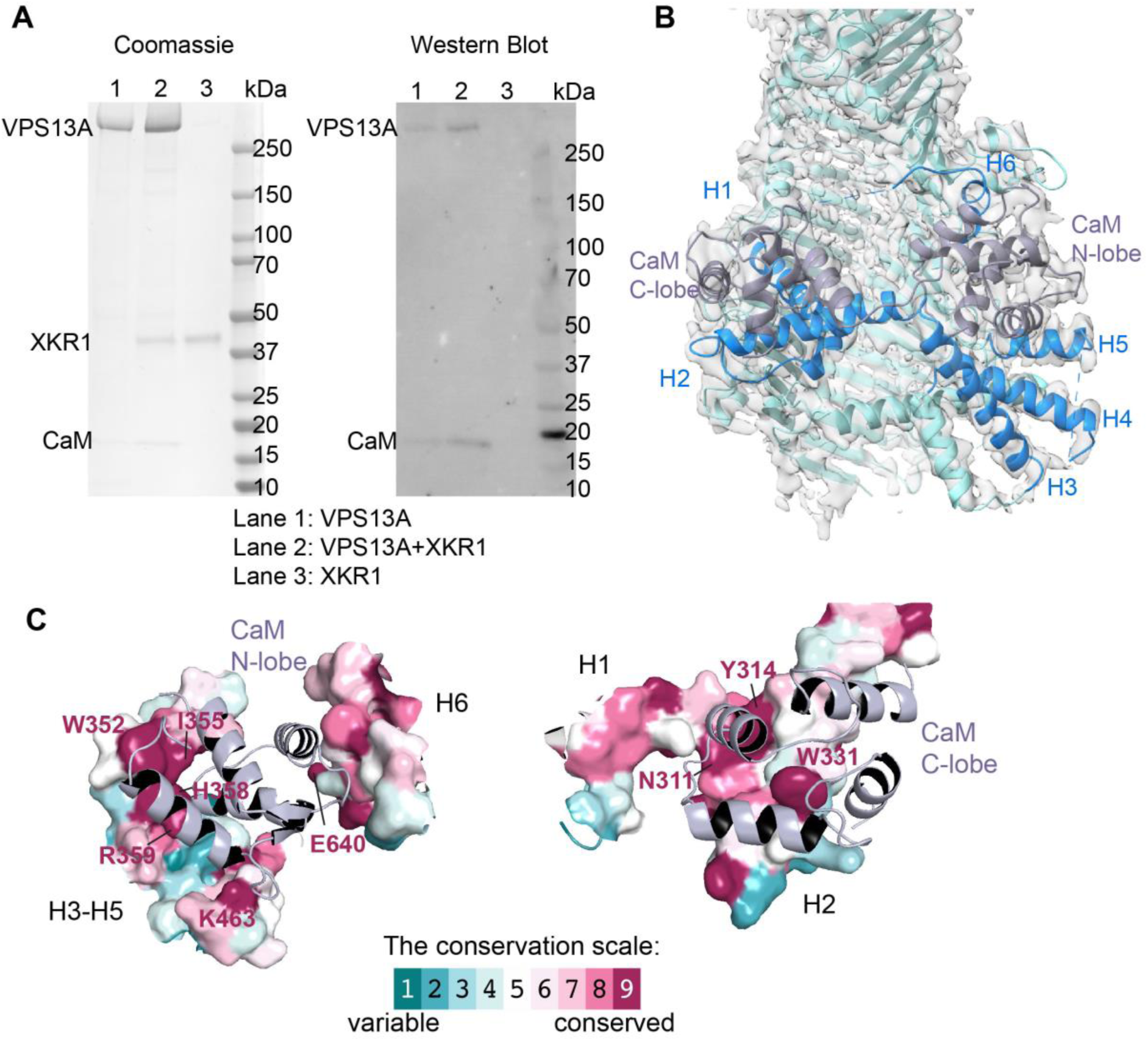
CaM binds to the N-terminus of VPS13A. **A**, CaM co-purifies with VPS13A, as validated by SDS-PAGE stained with Coomassie and western blot analysis using anti-CaM antibody. **B**, Locally refined cryo-EM map focused on the VPS13A-CaM region with the fitted atomic model. CaM is colored grey, the VPS13A helices interacting with CaM are colored blue, and the remainder of VPS13A is shown in light blue. **C**, Amino acids of VPS13A CaM binding helices are conserved. Conservation scores of residues interacting with the N-lobe and C-lobe of CaM were analyzed via Consurf (https://consurf.tau.ac.il/consurf_index.php). Only surfaces of VPS13A within 10 Å of CaM are shown. CaM has two domains, each comprising two EF-hands, which are helix-turn helix motifs with calcium binding sites in the loop between helices; and one or both domains of CaM can bind partner proteins in a calcium-sensitive manner. As observed for other VPS13s (Li et al., 2025), VPS13A engages with both lobes of CaM via a multi-helix segment that spans over the taco-shell bridge connecting the first and second RBGs. CaM contacts H292-312, H327-350, H352-374, H395-412, and H460-470 in VPS13A. Additionally, in VPS13A, the N-terminal lobe of CaM packs against loop elements of RBG 3 (residues 630-645 and 682-706). The total occluded surface area between VPS13A and CaM is large (5700 A^2^), indicative of a high affinity complex.

**Figure S2.**
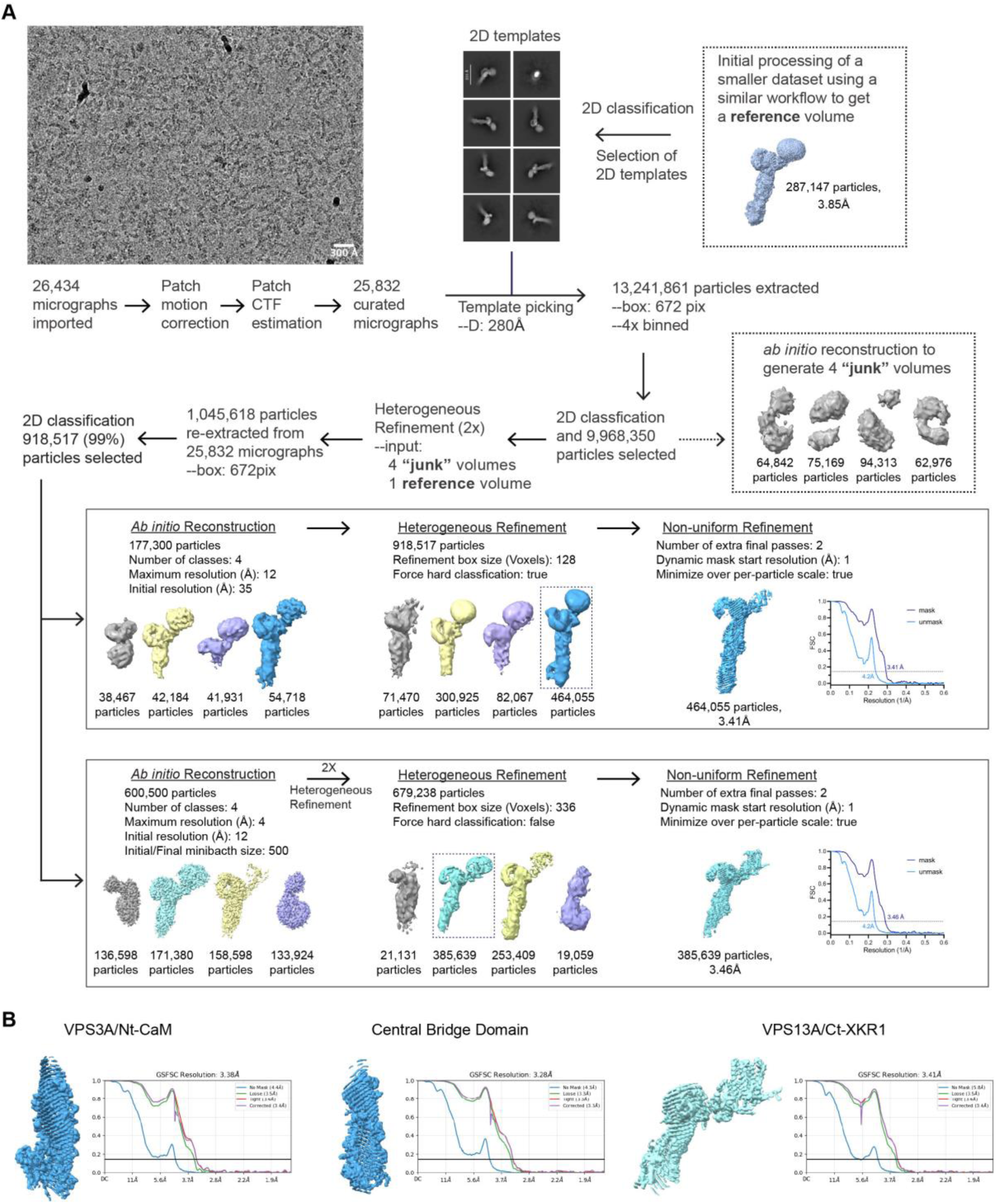
Cryo-EM data processing workflow for the VPS13A/CaM-XKR1 complex. **(A)** Data were processed in CryoSPARC (v4.6.2). A representative micrograph and 2D templates were shown. (**B)** Local refinement maps.

**Figure S3.**
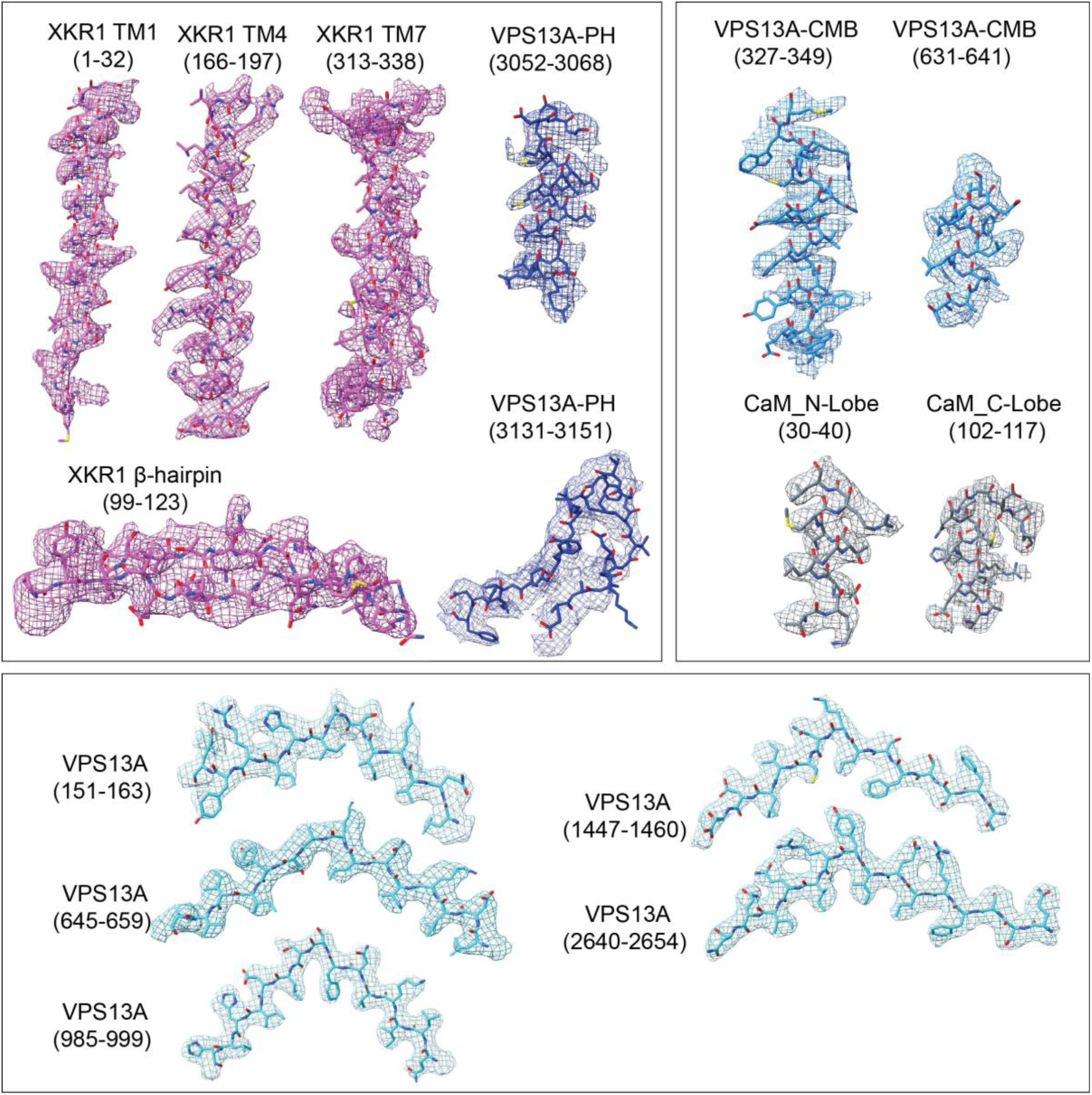
Selected atomic model of VPS13A/CaM-XKR1 in different regions of the density maps. The cryo-EM maps are shown as mesh. Color scheme is the same as that in Figure 1. CBM: calmodulin (CaM) binding motif.

**Figure S4.**
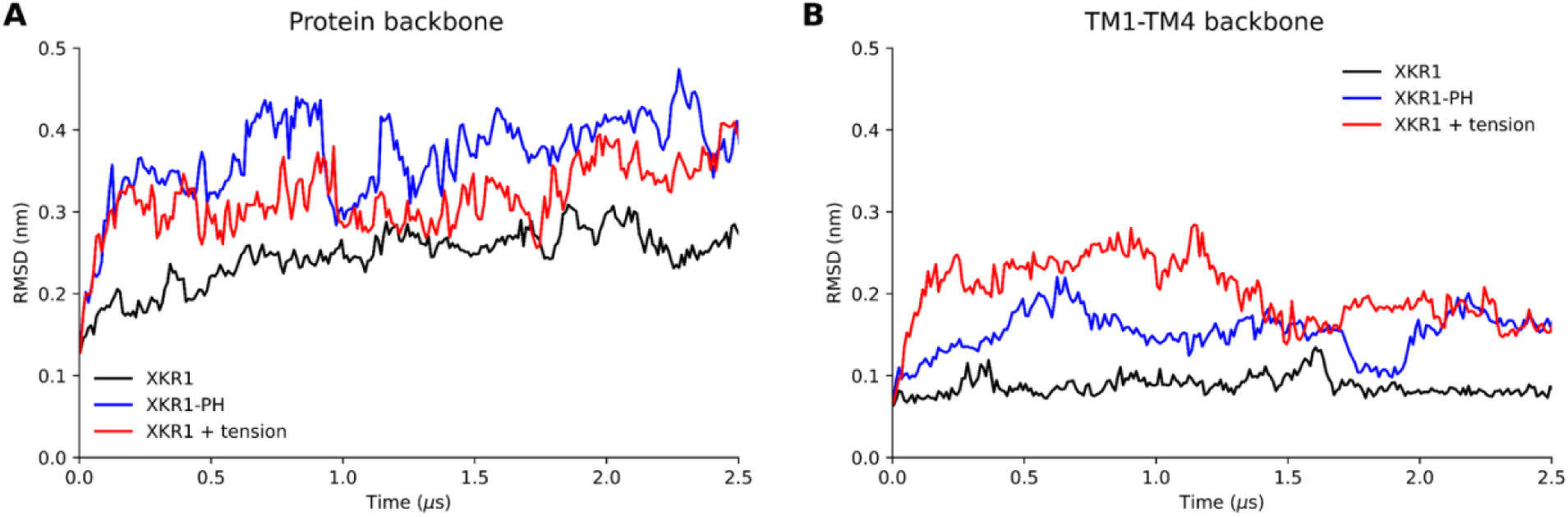
Time traces of the root mean squared deviation of the backbone for the XKR1 (A) and for the TM1-TM4 separation (B). Three conditions are shown, XKR1 alone (black), in presence of the PH domain (blue), and with membrane tension (red). The presence of the PH domain and the applied membrane tension led to an open conformation.

**Figure S5.**
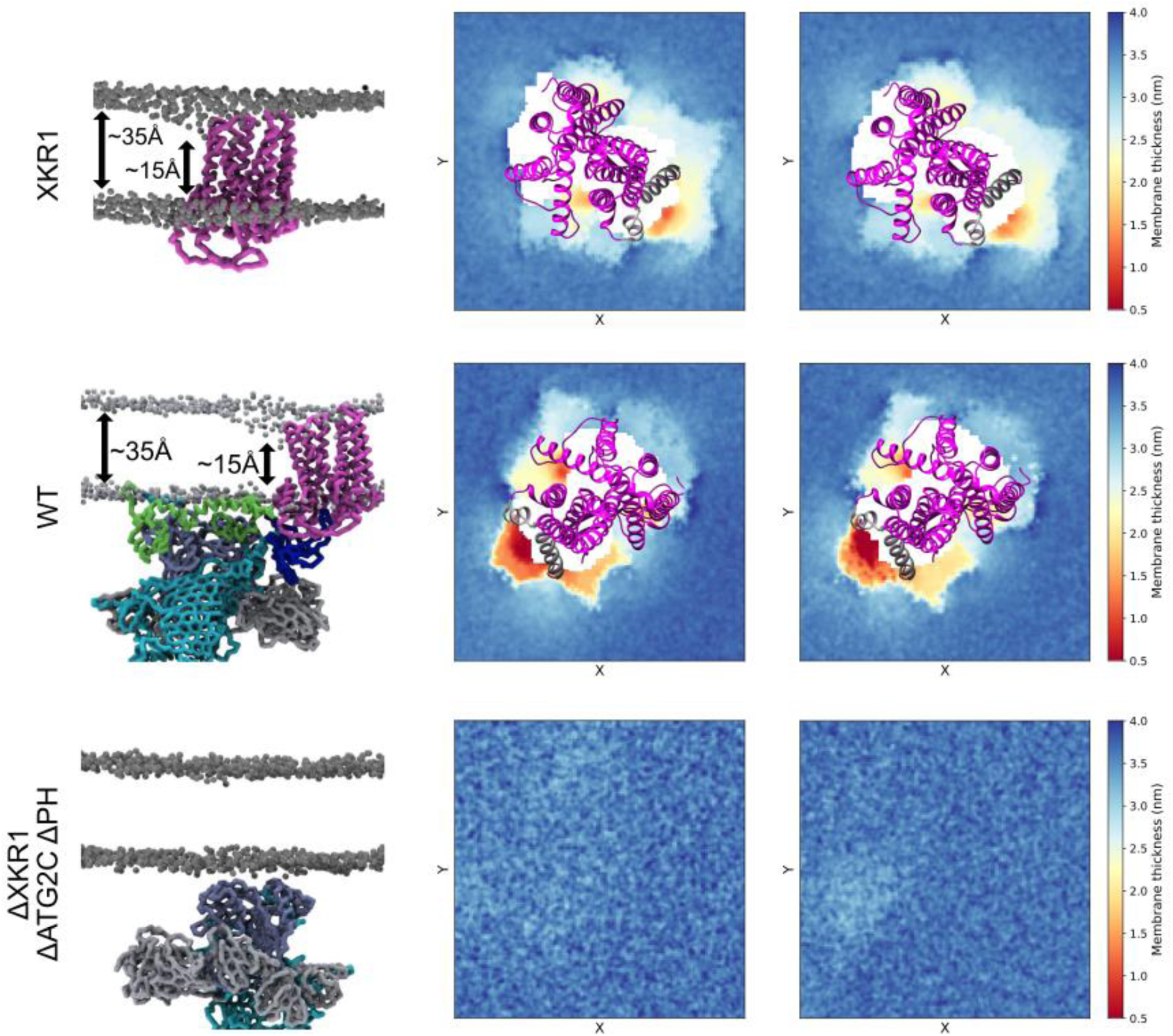
Membrane thickness in nm of the CG-MD simulations of XKR1, VPS13A/calmodulin-XKR1, and VPS13A (ΔATG2_C ΔPH)/calmodulin, with snapshots of the corresponding systems.

**Supplementary Figure S6.**
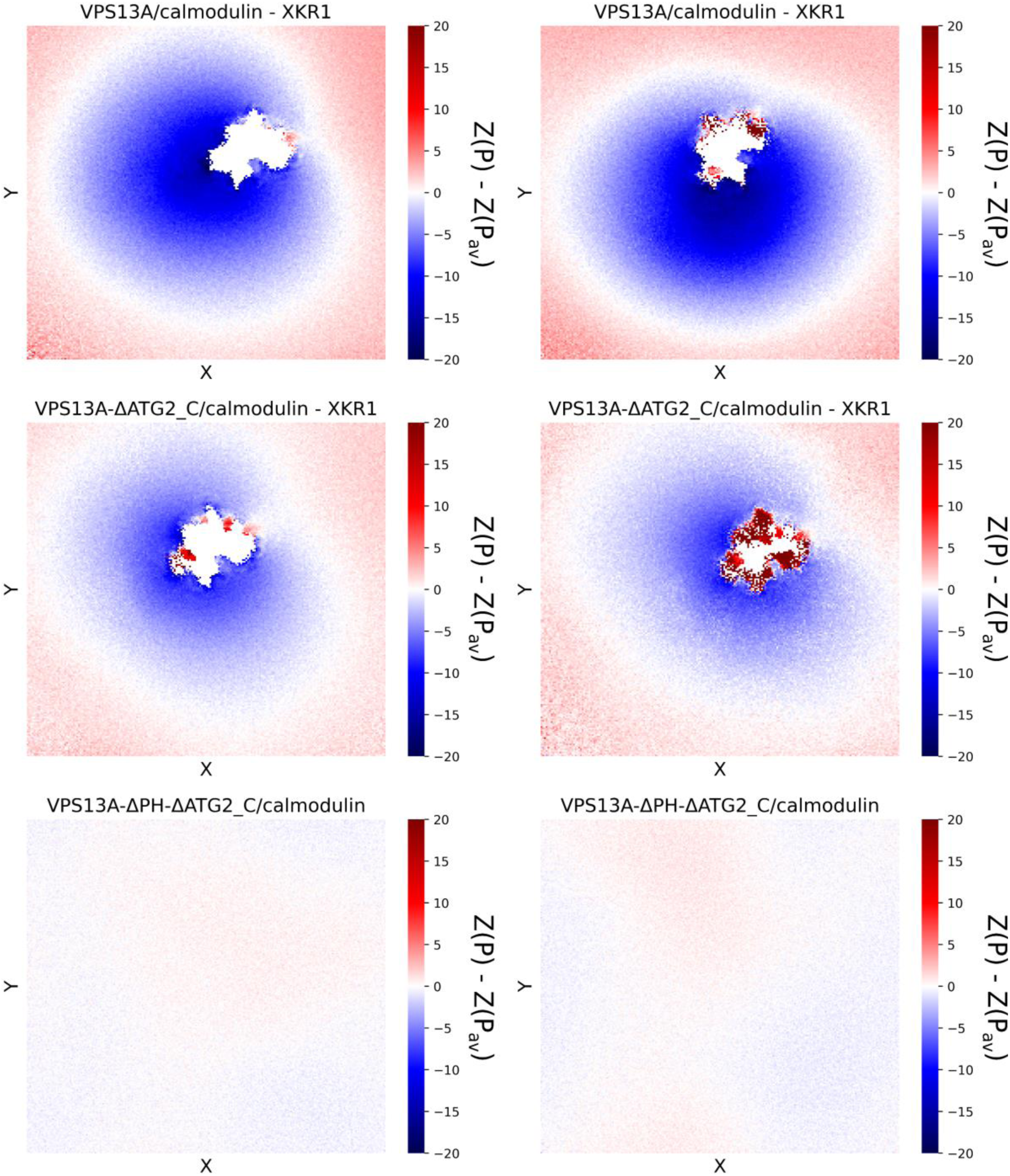
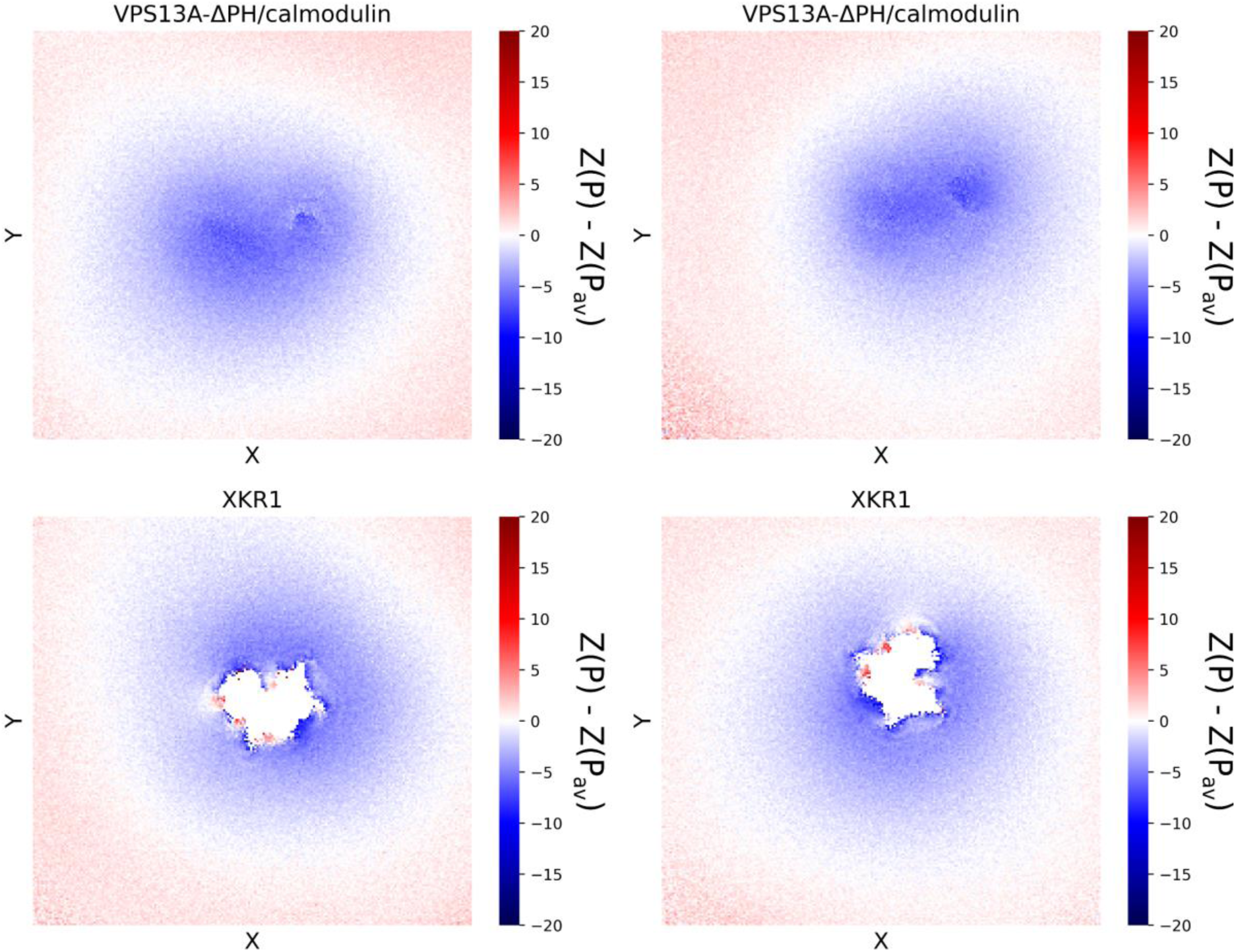
Grid x/y maps of the difference in Å between the z coordinate of the phosphate beads of the lipids in the cytosolic leaflet with respect to the average value, for the CG-MD simulations of the VPS13A/calmodulin-XKR1 complex and its mutants, as well as of XKR1 embedded in the membrane

**Figure S7.**
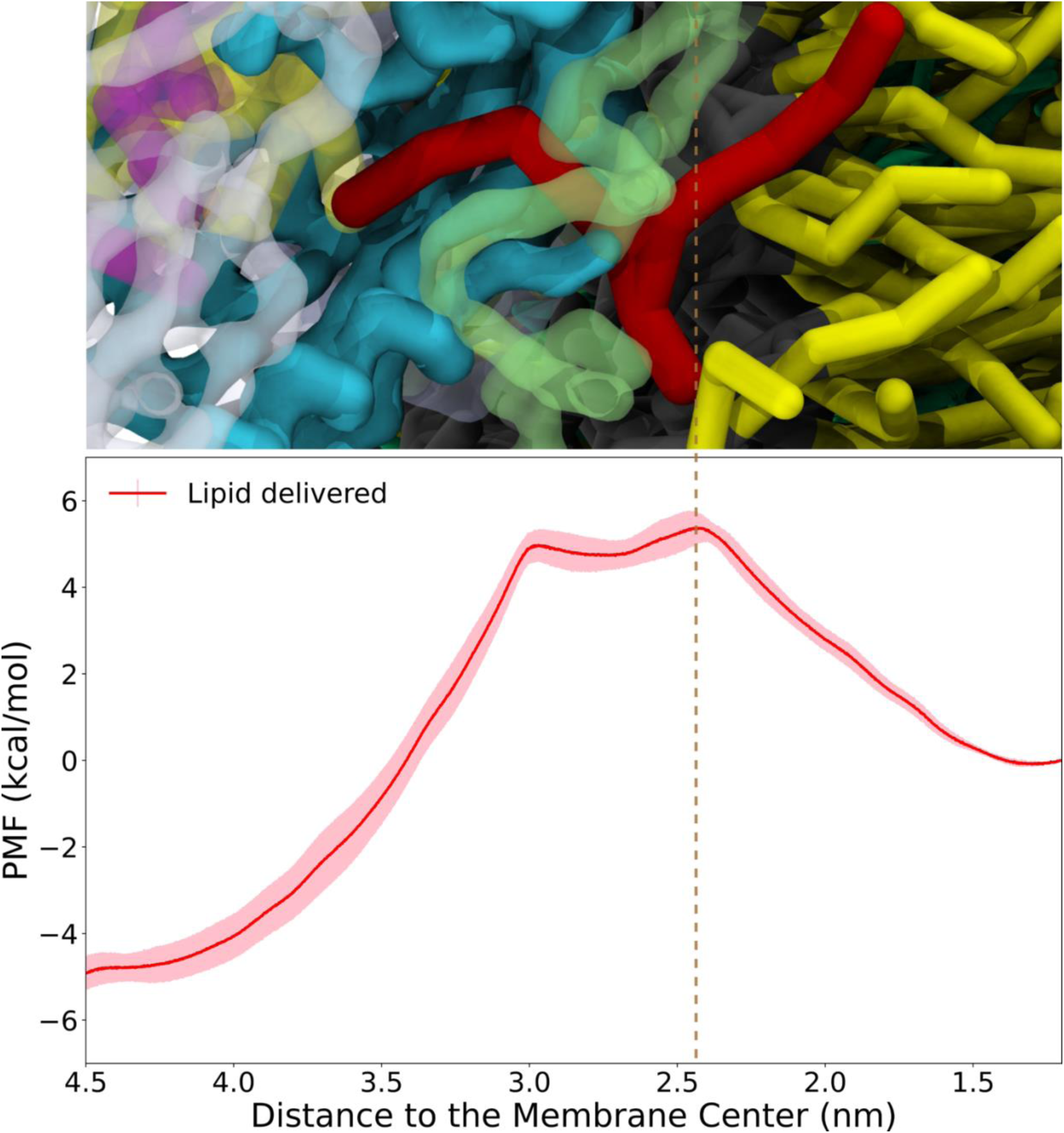
Potential of mean force (PMF) in kcal mol^-1^ of a lipid being delivered (red lipid in the snapshot) from VPS13A with 54 lipids placed inside its cavity, from the hydrophobic cavity (left) to the membrane (right) along the reaction coordinate. The vertical dotted line indicates the position of the membrane headgroups.

**Figure S8.**
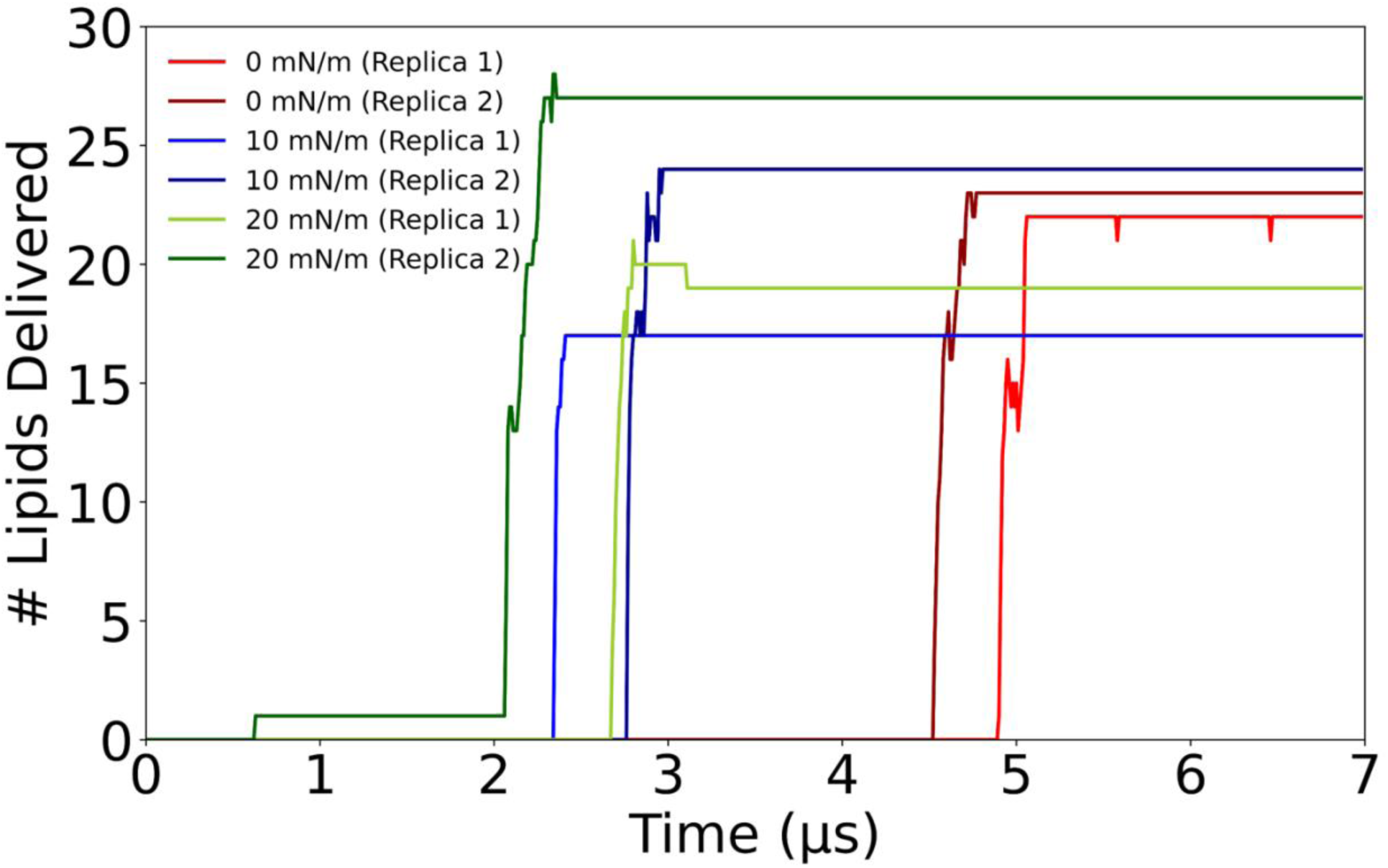
Time traces of the number of delivered lipids from VPS13A to the membrane in the CG MD simulations with 93 lipids initially placed inside the protein.

**Figure S9.**
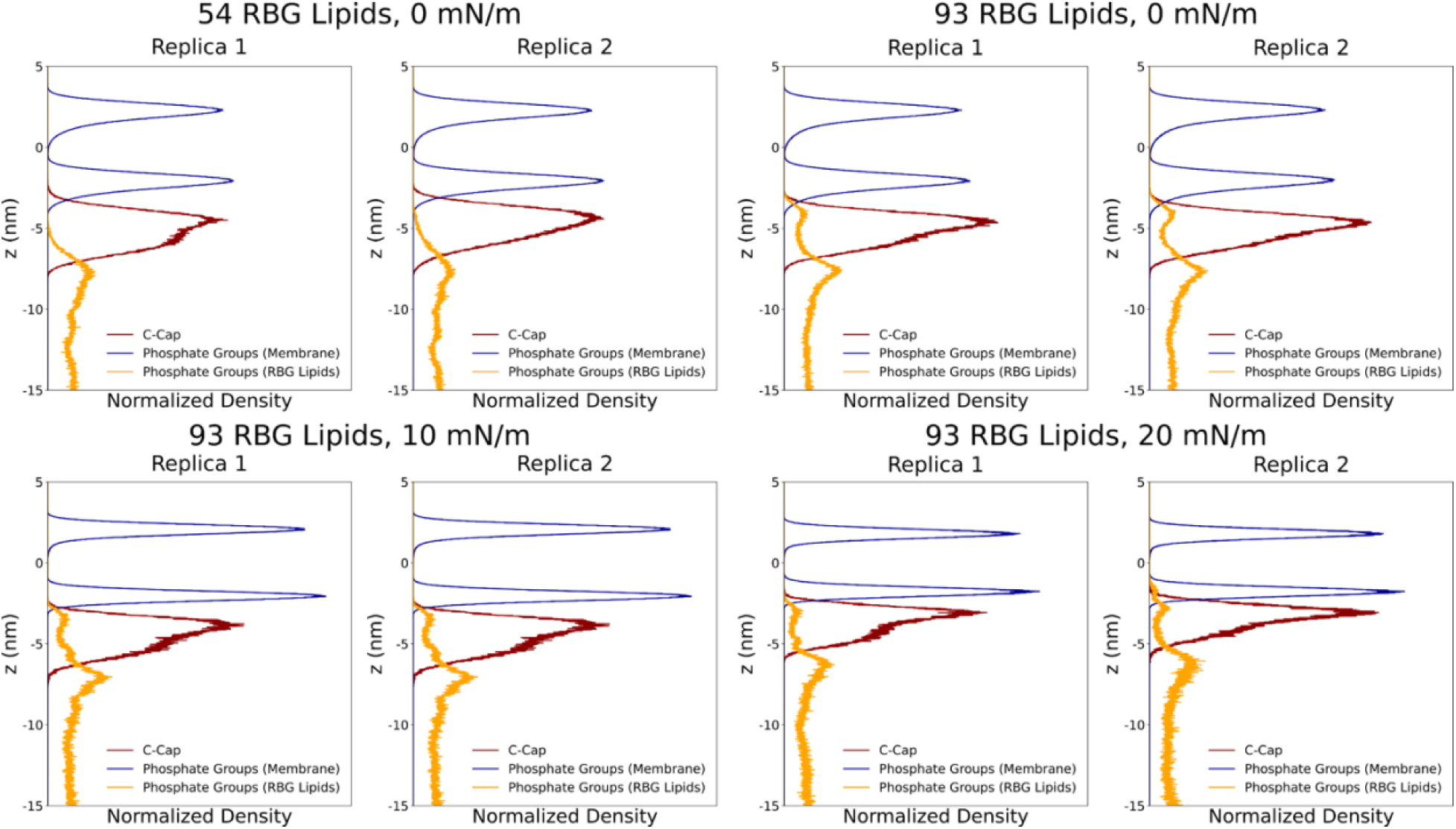
Normalized density profiles along z of the phosphate groups of the membrane lipids (in blue) and of the lipids initially placed inside the hydrophobic cavity of VPS13A (RBG lipids, in orange), as well as of the C-cap of VPS13A (in red) for the CG-MD simulations with either 54 or 93 RBG lipids, under different tension conditions. The profiles were computed across the simulation time before lipid delivery (if any).

**Supplementary Table S1.** Cryo-EM data collection, refinement and validation statistics.

|  | VPS13A/Nt-CaM | VPS13A Central Bridge domain | VPS13A/Ct-XKR1 |
| --- | --- | --- | --- |
|  | PDB 9YG4 | PDB 9YFW | PDB 9YG5 |
|  | EMD-72912 | EMD-72909 | EMD-72913 |
| **Data Collection and Processing** |  |  |  |
| Facility | Yale West Campus-Cryo-EM facility |  |  |
| Microscope | Titan Krios |  |  |
| Voltage (kV) | 300 |  |  |
| Camera | K3 |  |  |
| Magnification | 105,000 |  |  |
| Pixel Size (Å/px) | 0.832 |  |  |
| Total Electron Exposure (e-/Å<sup>2</sup>) | 50 |  |  |
| Defocus Range (µm) | -0.8 to -2.0 |  |  |
| Symmetry Imposed | C1 |  |  |
| Num of mics | 26434 |  |  |
| Initial Particles | 918,517 | 918,517 | 918,517 |
| Final Particles | 464,055 | 464,055 | 385,639 |
| Map Resolution (Å) | 3.38 | 3.28 | 3.41 |
| FSC threshold | 0.143 | 0.143 | 0.143 |
| **Refinement** |  |  |  |
| Initial models | AlphaFold | AlphaFold | AlphaFold |
| Model Resolution (Å) | 3.30 | 3.10 | 3.30 |
| FSC threshold | 0.143 | 0.143 | 0.143 |
| **Model Composition** |  |  |  |
| Non-hydrogen atoms | 6741 | 4543 | 8251 |
| Protein residues | 831 | 570 | 1020 |
| Ligands | 0 | 0 | 0 |
| **B factors (Å<sup>2</sup>)** |  |  |  |
| Protein | 122.17 | 69.91 | 107.64 |
| Ligands | - | - | - |
| **R.m.s deviation** |  |  |  |
| Bond length (Å) | 0.004 | 0.004 | 0.004 |
| Bond angles (°) | 1.044 | 0.648 | 1.000 |
| **Validation** |  |  |  |
| Molprobity score | 2.06 | 1.89 | 1.89 |
| Clashscore | 16.00 | 11.32 | 14.11 |
| Rotamer outliers (%) | 0.79 | 0.77 | 1.08 |
| **Ramachandran plot** |  |  |  |
| Outliers (%) | 0 | 0 | 0.00 |
| Allowed (%) | 5.13 | 4.64 | 3.59 |
| Favored (%) | 94.87 | 95.36 | 96.41 |

**Supplementary Table S2.**
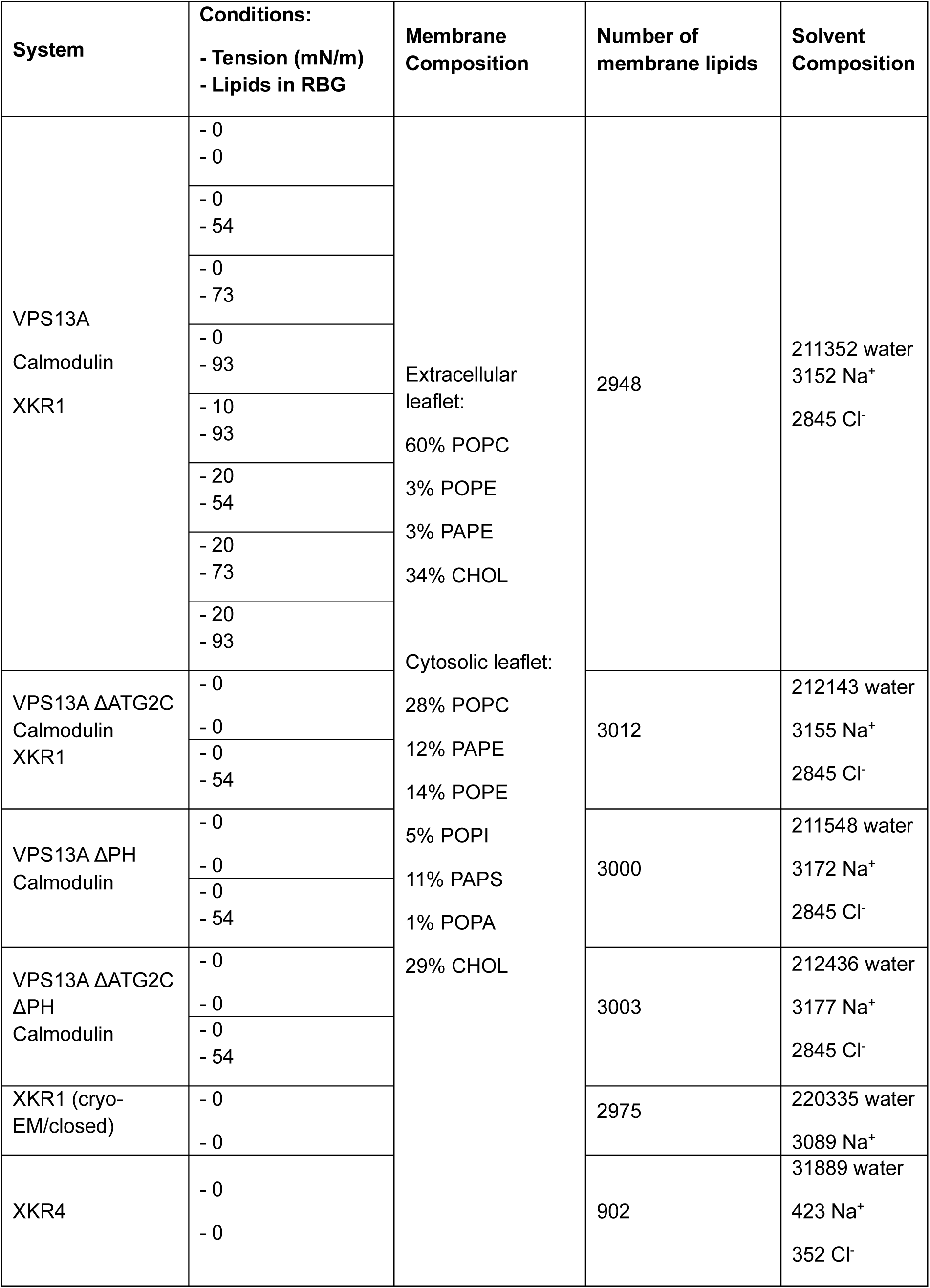
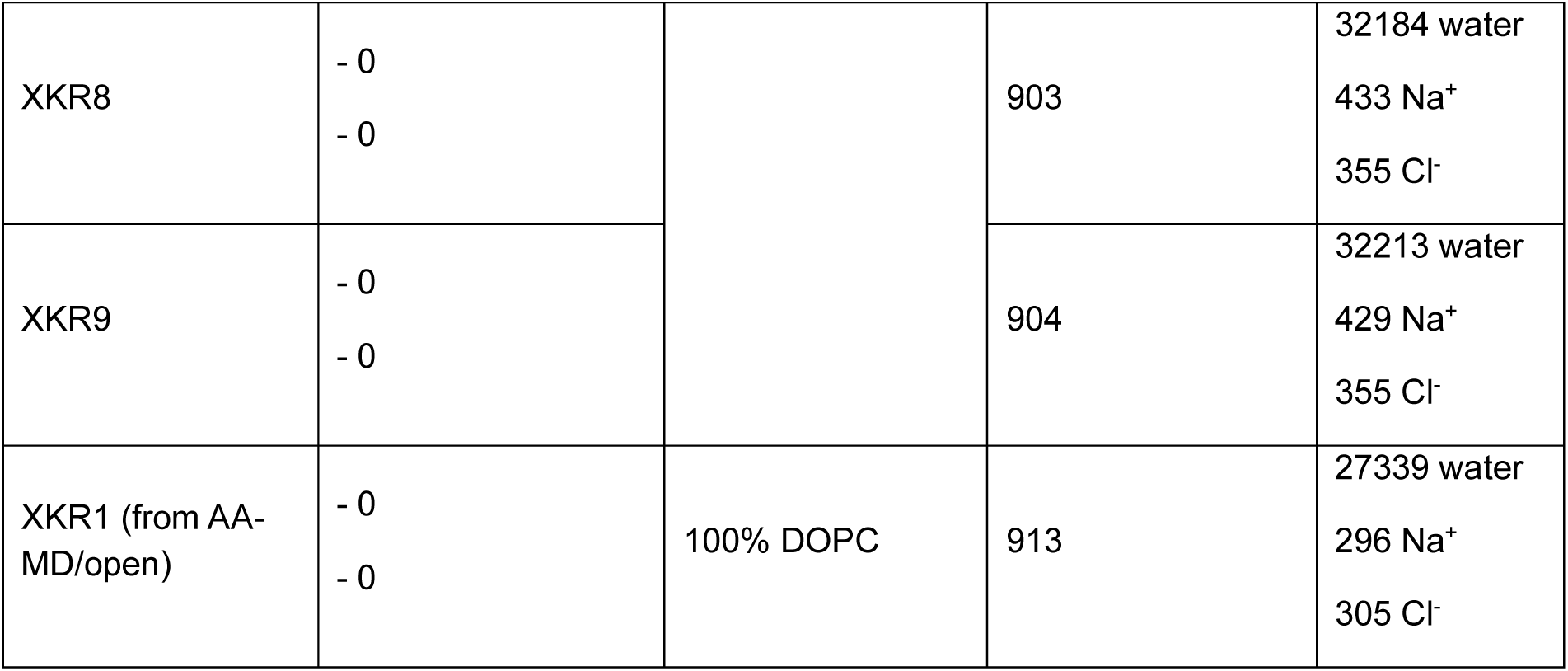
Tension conditions in mN/m, initial lipids inside the VPS13A hydrophobic cavity, membrane composition (in %) of each lipid along with the total number of lipids, and solvent composition for all the CG-MD protein-membrane systems simulated.

**Supplementary Table S3.**
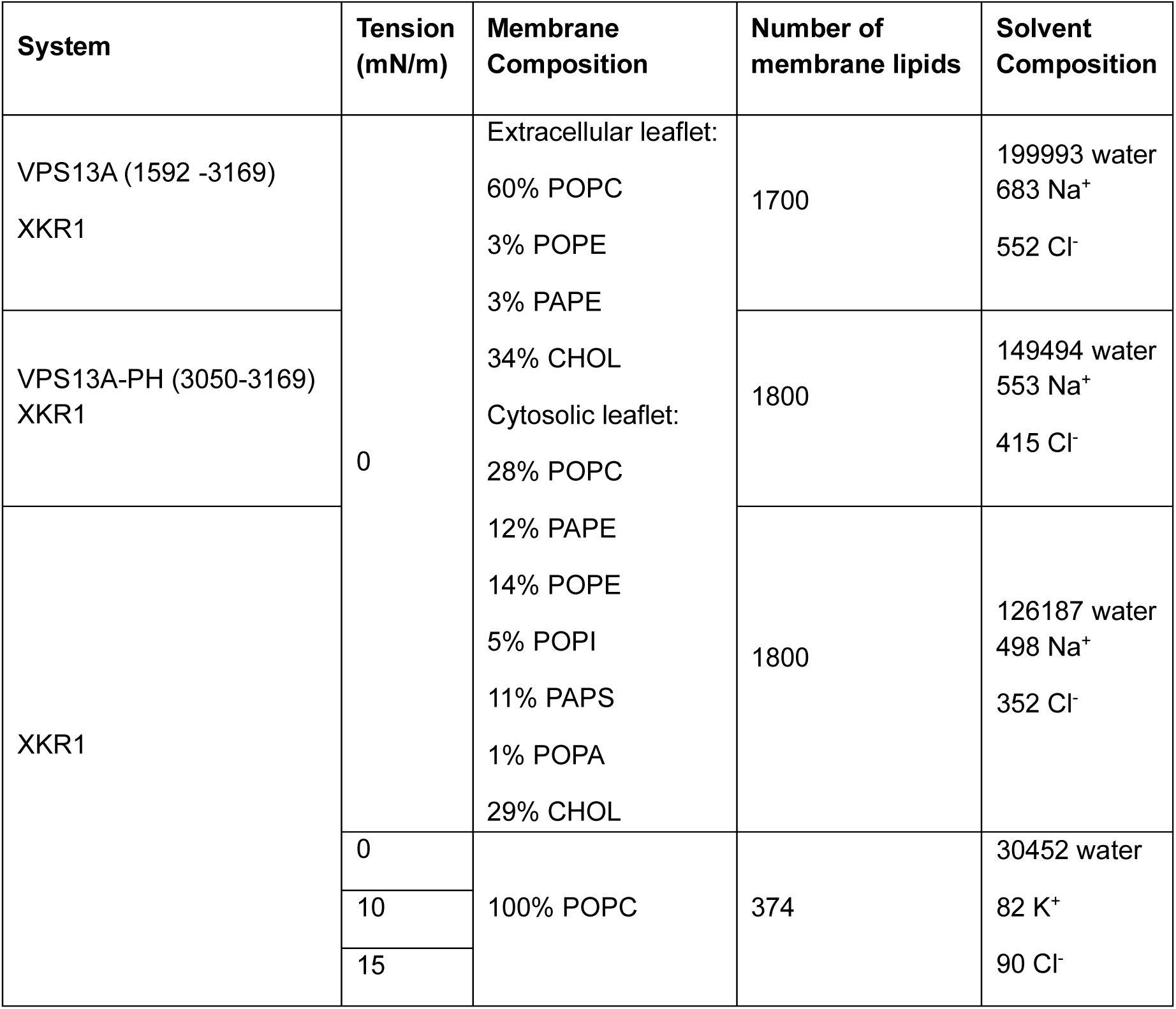
Tension conditions in mN/m, membrane composition (in %) of each lipid along with the total number of lipids, and solvent composition for all the AA-MD protein-membrane systems simulated.

**Supplementary Movie S1.** Delivery of lipids from the hydrophobic cavity of VPS13A in the CG-MD of VPS13A in complex with calmodulin and XKR1 and 93 lipids initially placed inside the hydrophobic cavity, without tension in the lipid membrane. Lipids initially placed in VPS13A are displayed with the lipid tails in yellow and the headgroup and backbone beads in purple, the headgroup of the membrane lipids is shown as black spheres.

**Supplementary Movie S2.** Delivery of lipids from the hydrophobic cavity of VPS13A in the CG-MD of VPS13A in complex with calmodulin and XKR1 and 93 lipids initially placed inside the hydrophobic cavity, with 20 mN/m of tension in the lipid membrane. Lipids initially placed in VPS13A are displayed with the lipid tails in yellow and the headgroup and backbone beads in purple, the headgroup of the membrane lipids is shown as black spheres.

## Notes

### Competing Interest Statement

The authors have declared no competing interest.

